# Butterfly eyespots evolved via co-option of the antennal gene-regulatory network

**DOI:** 10.1101/2021.03.01.429915

**Authors:** Suriya Narayanan Murugesan, Heidi Connahs, Yuji Matsuoka, Mainak das Gupta, Manizah Huq, V Gowri, Sarah Monroe, Kevin D. Deem, Thomas Werner, Yoshinori Tomoyasu, Antónia Monteiro

**Affiliations:** Department of Biological Sciences, National University of Singapore; Department of Biology, Miami University, USA; Department of Biological Sciences, Michigan Technological University, USA; Science Division, Yale-NUS College, Singapore

## Abstract

Butterfly eyespots are beautiful novel traits with an unknown developmental origin. Here we show that eyespots likely originated via co-option of the antennal gene-regulatory network (GRN) to novel locations on the wing. Using comparative transcriptome analysis, we show that eyespots cluster with antennae relative to multiple other tissues. Furthermore, three genes essential for eyespot development (*Distal-less* (*Dll*), *spalt* (*sal*), and *Antennapedia* (*Antp*)) share similar regulatory connections as those observed in the antennal GRN. CRISPR knockout of *cis*-regulatory elements (CREs) for *Dll* and *sal* led to the loss of eyespots and antennae, and also legs and wings, demonstrating that these CREs are highly pleiotropic. We conclude that eyespots likely re-used the ancient antennal GRN, a network previously implicated also in the development of legs and wings.

## Main text

Although the hypothesis of GRN co-option is a plausible model to explain the origin of morphological novelties (*1*), there has been limited empirical evidence to show that this mechanism led to the origin of any novel trait. Several hypotheses have been proposed for the origin of butterfly eyespots, a novel morphological trait. These include GRN co-option from the leg (*2*), embryo segmentation (*3*), wing margin (*4*) and wound healing (*5*). These hypotheses for eyespot GRN origins all rely on similarities of expression of just a few candidate genes observed in eyespots and in the proposed ancestral gene network. To test whether co-option of any of these networks underlies eyespot origins, we focused on the nymphalid butterfly *Bicyclus anynana*, which has served as a model for studying eyespot development (*6*). Using RNA-sequencing (RNA-seq), we examined and compared the larger collection of genes expressed in a forewing eyespot of *B. anynana* with those expressed in these proposed candidate ancestral traits. Additionally, we examined a few other traits, including larval head horns and prolegs, and also pupal eyes and antennae (Fig. 1A).

**Fig. 1:**
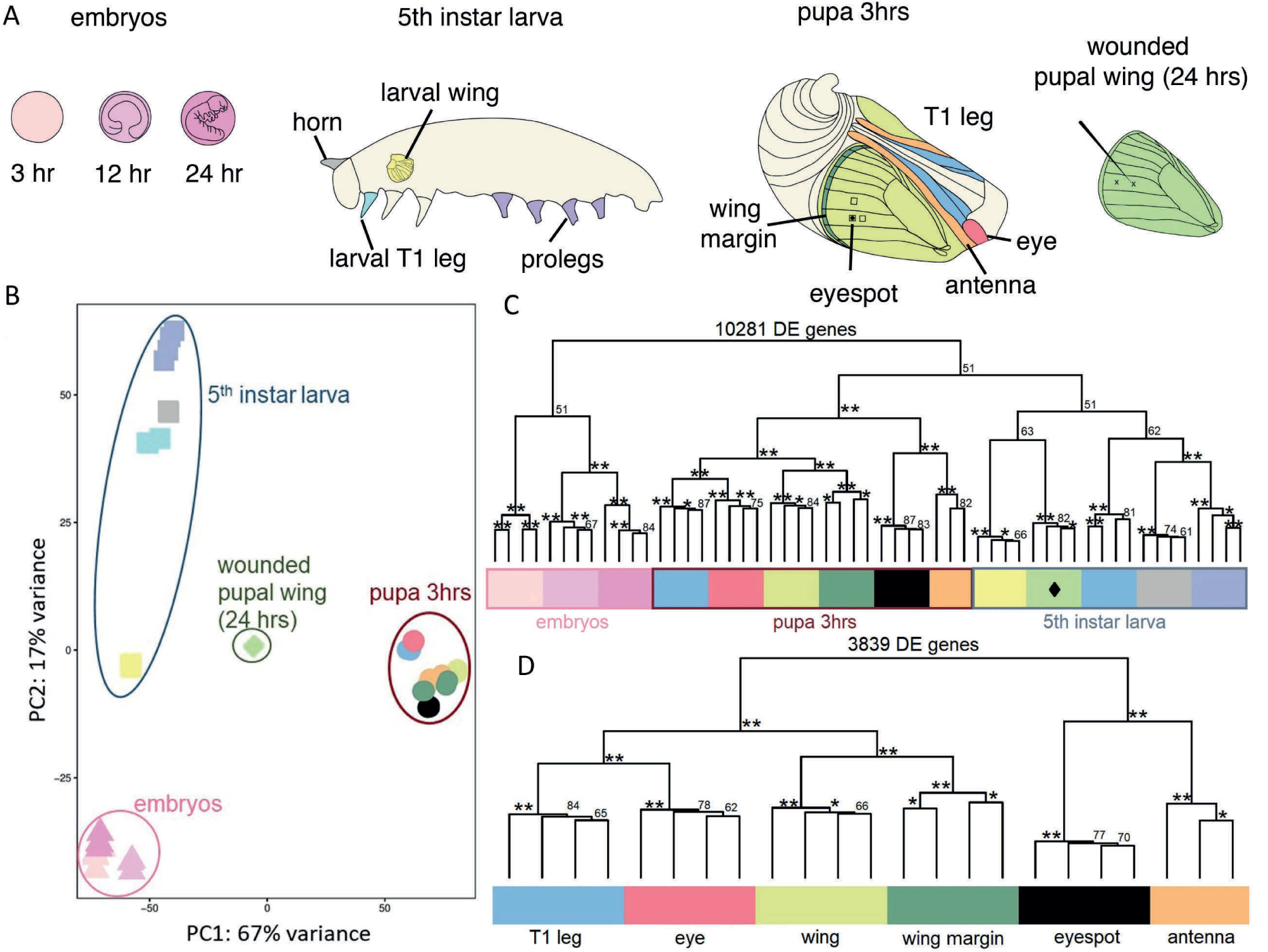
Tissues used for RNA-seq analysis and character tree constructed using the differentially expressed (DE) genes. (A). We used 16 tissue groups from three separate developmental stages of *B anynana* for RNA extractions. Embryos at 3 hrs, 12 hrs and 24 hrs after egg laying. Larval forewings, T1-legs, horns and prolegs. Pupal antenna, T1-leg, forewing, eye, wing margin, eyespot, and two eyespot control tissues all dissected at 3hrs after pupation, and a wounded wing dissected at 24hrs after pupation. (B). PCA using 10281 DE genes obtained from pairwise comparisons between different tissues. Tissues are clustering according to their developmental stages. (C). Character tree constructed using 10281 DE genes showing eyespot tissue clustering with antenna tissue first, and next with tissues from same developmental stage, except for a 24 hrs wounded wing (◆), which clustered with larval wing tissue (D). Character tree constructed using 3839 DE genes from 3hr pupal stage showing eyespot tissue clusters with antenna tissue, which together form an outgroup to the rest of the samples. ** - 100 unbiased (AU) p-value; *- 90-99 unbiased (AU) p-value; ◆ - wounded pupal wing (24 hrs)

### The transcriptome profile of eyespots and antennae cluster together

We first examined which of the sampled tissues shared the most similar gene expression profile to eyespot tissue, as these should cluster closer together (*7*). Pairwise differential expression (DE) analysis using DESeq2 (*8*) identified 10,281 DE genes (logFC ≥ |2| and padj ≤ 0.001) among all tissues sampled. Hierarchical clustering of tissues, using DE genes, resulted in eyespots clustering with antennae (Fig. 1B), but tissues were also clustering according to developmental stage (Fig. 1B, 1C). To circumvent the strong developmental stage signal, we reanalyzed DE genes solely from 3-h-old pupae, when the eyespot tissue was dissected. We found 3,839 DE genes between the tissues, with eyespots clustering with antennae, and both forming an outgroup to the remaining tissues with a high approximately unbiased (AU) *P*-value (*9*) (Fig. 1D).

To more narrowly identify the subset of genes associated with eyespot development and to examine similarities in their expression profile with our candidate tissues, we next compared the transcriptome of dissected eyespot tissue with adjoining control tissue in the same wing sector (Fig. 1A), as done by a previous study (*10*). This previous study identified 183 genes differentially expressed in eyespots relative to sectors of the wing without eyespots. Our new DE analysis between eyespot and control wing tissues identified 652 eyespot-specific DE genes with 370 being up-regulated, which included *sal*, and 282 down-regulated in eyespots (Fig. S1, S2, Spreadsheet S1). We mapped the published 183 eyespot DE genes, which included *Dll* and *Antp*, to the current assembled transcriptome. After removing multi-mapped genes, we retained 144 genes from the published study for further analysis (Spreadsheet S1). When hierarchical clustering was performed, using either the newly identified 652 genes, the 144 genes previously identified, or both datasets combined, we found that the eyespot transcriptome always clustered with antennae with strong support AU *P*-value for the clade. This clustering persisted with just the 370 up-regulated genes (Fig. S3A, E, F).

Given the importance of transcription factors (TFs) in development and in establishing GRNs, we used 336 genes annotated as having “DNA-binding transcription factor activity (GO:0003700)” and “transcription factor binding (GO:0008134)” in a separate analysis, which showed eyespots again clustering with antennae (Fig. S3B). Annotation and gene enrichment for the DE genes (3, 839) between the 3-h-pupal stage tissues showed a strong enrichment in animal organ morphogenesis (GO:0009887) and anatomical structure formation (GO:2000026) (Fig. S4). Performing the clustering analysis using genes from these two groups (GO:0009887 and GO:2000026), in two separate analyses, reproduced the same results as the full gene set, indicating that these morphogenesis genes show similar expression profiles in both eyespots and antennae (Fig. S3C, 3D).

These analyses showed that eyespots and antennae form an outgroup to the other tissues, including legs, which are considered serial homologs to antennae. However, eyespots express a key selector gene, *Antp* which is known to give legs their unique identity and differentiate them from antennae. Antp protein is known to positively regulate *Dll* and repress *sal* in the leg disc of *Drosophila* (*11, 12*), whereas in the antennae, in the absence of Antp, Dll activates *sal* (*13*). Comparative data across 23 butterfly species suggested that eyespots originated without Antp protein expression, and that *Antp* was recruited later to the eyespot GRN in at least two separate lineages, including in the ancestors of *B. anynana* (*14*). We therefore reasoned that if eyespots are co-opted antennae, rather than co-opted legs, the regulatory interactions between *Dll, Sal*, and *Antp* in eyespots should resemble those in insect antennae but not those in legs, and that the regulatory interactions between *Antp* and the other two genes should be novel and not homologous.

### Function of *sal* and regulatory interactions between *Dll*, *sal*, and *Antp* in eyespots

Before establishing regulatory interactions between the three genes, we first obtained missing functional data for one of these genes, *sal,* lacking for *B. anynana*. Mutations for *Dll* and *Antp* were previously shown to remove eyespots, pointing to these genes as necessary for eyespot development (*6, 15*). We disrupted the function of *sal*, using CRISPR with a single guide RNA (sgRNA) targeting exon 2 (Fig. S5). *sal* crispants (mosaic mutants) showed a range of phenotypes, from missing eyespots (Fig. 2B and 2D, Fig. S6) to altered chevron patterns on the wing margin and the central symmetry system bands running the length of each wing (Fig. 2B), all mapping to patterns of *sal* expression in larval and pupal wings (Fig. 2H, 2K) (*5, 16*).

**Fig. 2.**
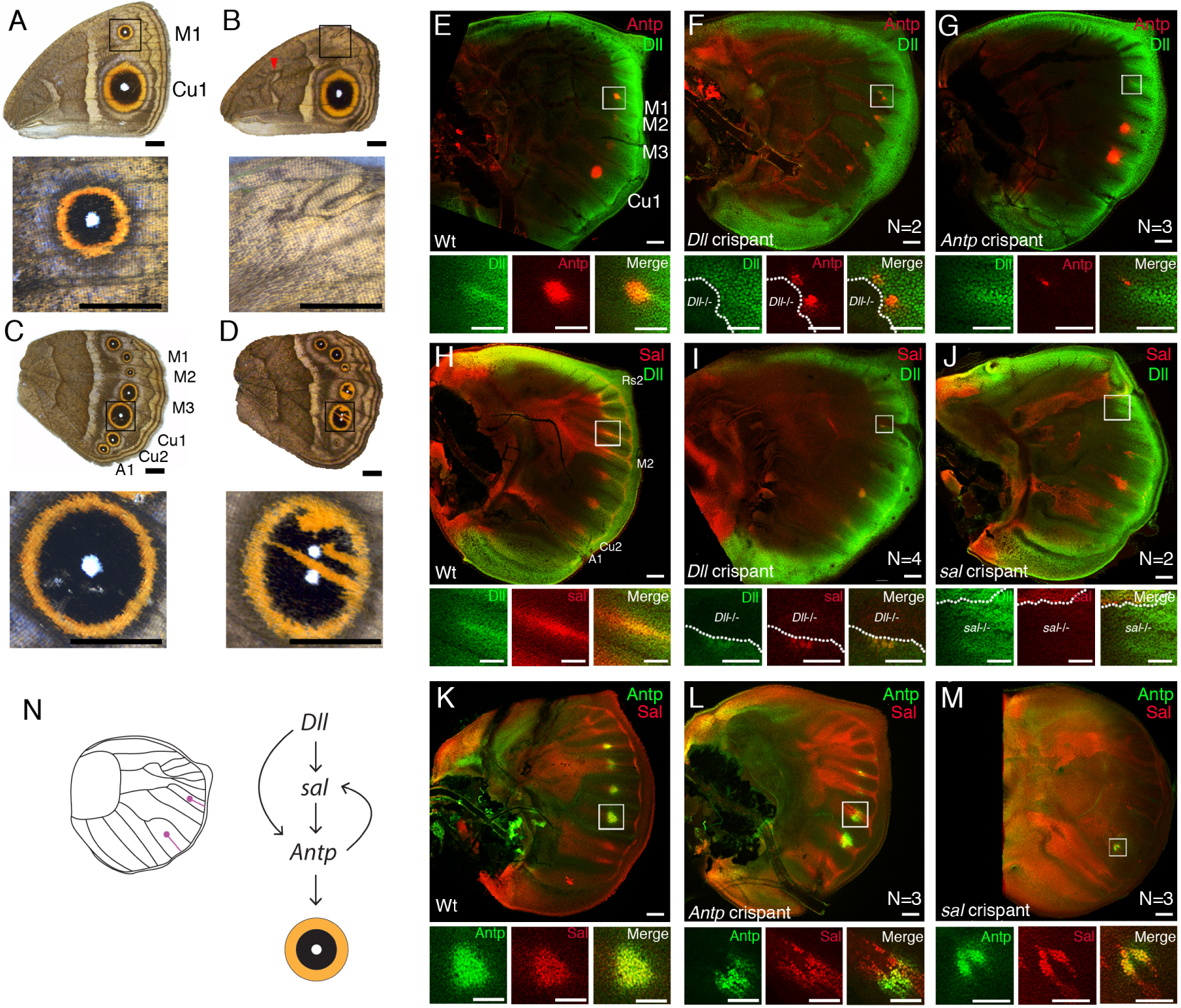
Function of *sal* and regulatory interactions between *Dll*, *sal*, and *Antp* inferred with CRISPR and immunohistochemistry. (A). Wt female forewing. (B) *sal* crispant female forewing (C) Wt female hindwing. (D) *sal* crispant female hindwing (E) Expression pattern of Dll and Antp proteins in Wt forewing. (F) Expression pattern of Dll and Antp proteins in *Dll* crispant forewing. (G) Expression pattern of Dll and Antp proteins in an *Antp* crispant forewing. (H) Expression pattern of Dll and Sal proteins in Wt forewing. (I) Expression pattern of Dll and Sal proteins in *Dll* crispant forewing. (J) Expression pattern of Dll and Sal proteins in *sal* crispant forewing. (K) Expression pattern of Sal and Antp proteins in Wt forewing. (L) Expression pattern of Sal and Antp proteins in *Antp* crispant forewing. (M) Expression pattern of Sal and Antp proteins in *sal* crispant forewing. White square regions were highly magnified. (N) Schematic diagram of genetic interaction among *Dll*, *sal*, and *Antp* in the eyespot region of a developing forewing. Scale bars in A-D: 5 mm for whole wings and wing details. Scale bars in E-M: 100 μm in low and 50 μm, in high magnification

Our data confirmed phenotypes previously shown in *J. coenia* (*17*). However, two novel and striking phenotypes were the splitting of eyespot centers into two smaller centers (Fig. 2D, Fig. S6) and the partial loss of black scales in the eyespot and their replacement with orange scales (Fig. 2D), resembling the “goldeneye” phenotype (*18*). Taken together, these results confirm that *sal* is necessary for the development of eyespots and also for the development of black scales.

To test the regulatory hierarchy between these three eyespot-essential genes, we knocked out each gene in turn, using CRISPR-Cas9, and reared the mosaic individuals until the late 5^th^ instar for larval wing dissections. We performed immunohistochemistry on these wings with antibodies against the protein of the targeted gene and against the other two proteins. We first examined the interaction of *Dll* with *Antp*. In wild-type (wt) wings, Dll protein is expressed along the wing margin and in finger-like patterns, spreading from the wing margin to the future eyespot centers (Fig. 2E), whereas Antp protein is initially expressed in the center of four putative eyespots (from M1 to Cu1) (*19*). In a *Dll* crispant forewing, Antp protein expression was affected in *Dll* null cells (Fig. 2F, Fig. S7), whereas Dll protein expression was not affected in *Antp* null cells in an *Antp* crispant (Fig. 2G, Fig. S8). These results suggest that *Dll* is upstream of *Antp* in eyespot development. We next examined the interaction of *Dll* with *sal.* In wt wings, Sal protein is broadly expressed along several wing sectors, connected to its role in vein patterning (*16*), and also expressed in nine potential eyespot centers (Fig. 2H and 2K). In *Dll* crispants, Sal expression was lost in *Dll* null clones in the eyespot centers (Fig. 2I, Fig. S9), but Dll protein expression was not affected in *sal* null clones in *sal* crispants (Fig. 2J, Fig. S10). These results suggest that *Dll* is also upstream of *sal* in eyespots. Finally, we examined the interaction between *Antp* and *sal*. In *Antp* crispants, Sal protein expression is missing from *Antp* null cells (Fig. 2L, Fig. S11). Furthermore, Antp protein expression is missing from *sal* null cells in *sal* crispants (Fig. 2M, Fig. S12). Taken together, *Dll* is up-regulating both *Antp* and *sal*, and *Antp* and *sal* are up-regulating each other’s expression in forewing eyespots (Fig. 2N).

### Regulatory connections between *Dll* and *sal* in eyespot development are similar to those in the antennae of flies

We next examined whether the appendage expression and regulatory connections between these three genes of *B. anynana* matched those known in fly leg and antennal development. In flies, Dll protein is expressed in both appendages (*20*), whereas Sal is only expressed in antennae and Antp only in legs of flies (*13*). In *B. anynana* we observed similar expression profiles in antennae and thoracic legs of pupae (Fig. S13-S14). Dll is necessary for *sal* expression in antennae of flies (*13*), as also observed in *B. anynana* eyespots (Fig. 2l). Antp, however, negatively regulates *sal* expression in fly legs (*12*), which differs from the regulation observed in eyespots, where *Antp* and *sal* up-regulate each other (Fig. 2N). The genetic interaction of *Antp* and *Dll* during leg development in *Drosophila* is stage-dependent. At the stage when leg primordia are formed, Antp positively regulates *Dll* expression in the thoracic leg bud (*11*), but when leg segments are being formed, Dll negatively regulates *Antp* in the distal leg elements (*21*). These regulatory interactions between *Dll* and *Antp* in leg development are distinct from the regulatory interaction observed in eyespots (Fig. 2N). Taken together, these data suggest that the regulatory interactions between *Dll* and *sal* in eyespots are likely homologous to those in the insect antenna GRN. *Antp* established a novel regulatory interaction to these two genes in eyespots, distinct from those found in the leg GRN of *Drosophila.* This supports the later and independent addition of *Antp* to the eyespot GRN in two separate lineages of butterflies, as proposed by Oliver et al. (2012) (*14*).

### Two pleiotropic CREs reveal a shared network between eyespots, antennae, and other traits

Evidence of GRN co-option is bolstered by the identification of shared *cis*-regulatory elements (CREs) driving the expression of genes common to both the ancestral and the novel trait (eyespots). To identify putative CREs specific to wing tissue with eyespots, we used Formaldehyde-Assisted Isolation of Regulatory Elements using sequencing (FAIRE-seq) to identify the open chromatin profile around *Dll* in forewing and hindwing pupal tissues of *B. anynana.* We produced separate libraries from the proximal and distal regions of the wing. Mapping of FAIRE-seq reads from each wing region to a previously published *Dll* BAC (scaffold length of 230 kb) revealed 18 regions of open chromatin across this scaffold, representing candidate CREs (Fig. 3A). A BLAST search of each candidate CRE against the *B. anynana* genome revealed that most of these regions contained repetitive elements.

**Fig. 3.**
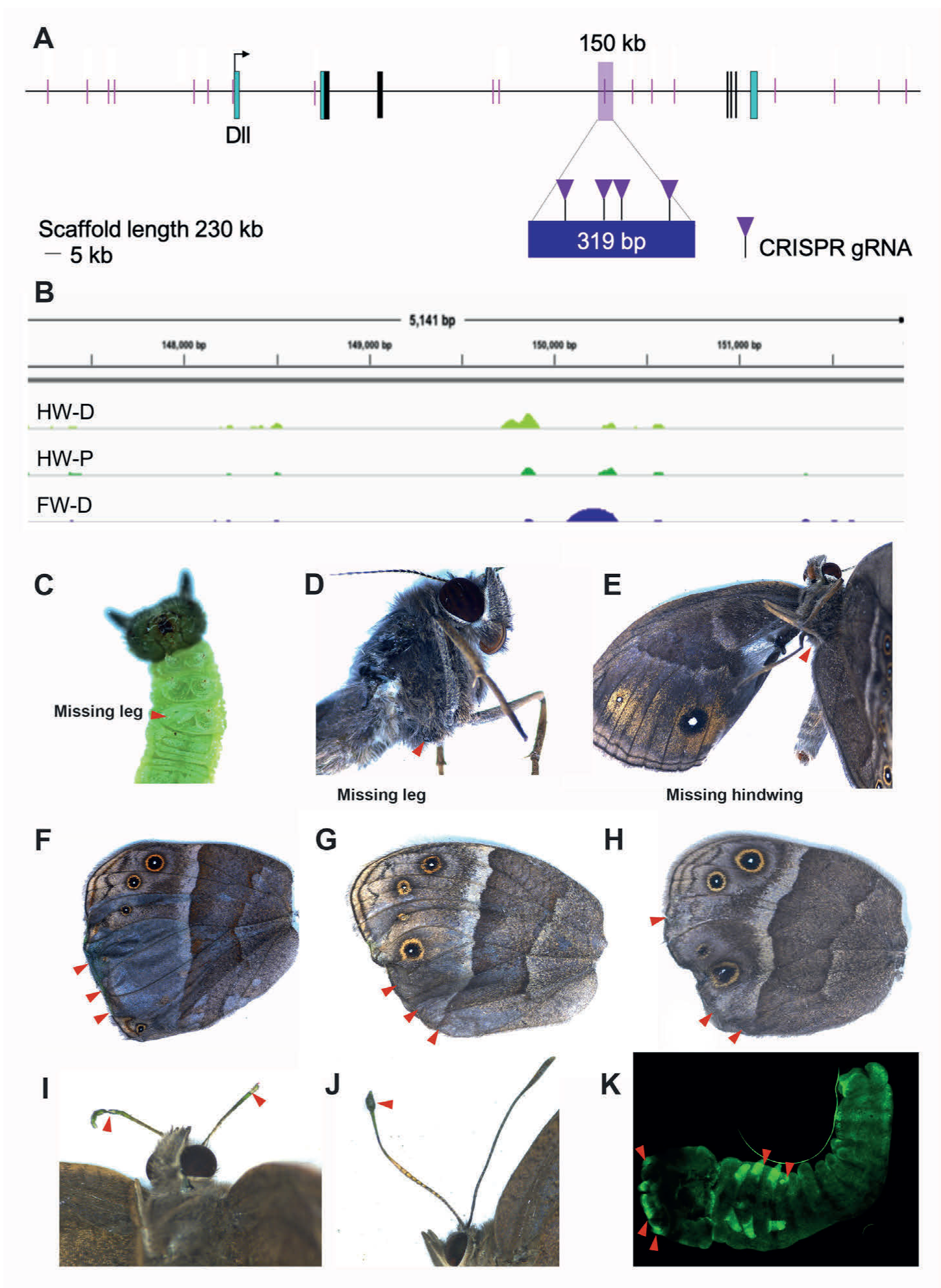
Multiple traits are affected by disruptions of a single *Distal-less* CRE. (A) *B. anynana Distal-less* BAC visualized using IGV showing all 18 FAIRE-seq open chromatin regions at 24 hours post-pupation (short red lines). First exon (UTR in blue) shows open chromatin region (highlighted by a short red line) at position 54 kb at the transcriptional start site of *Distal-less*. The CRE at position 150 kb (*Dll319*; highlighted with a pink bar) is open in *B. anynana* forewing and was targeted with CRISPR. Four RNA guides were used simultaneously to target this region. (B) FAIRE-seq results showing an open region of chromatin in the distal forewing (FW-D) at position 150 kb on the *Distal-less* BAC (blue peak). (C-E) crispant phenotypes from the same individual: with a missing thoracic leg as a caterpillar, and same missing thoracic leg and also missing hindwing as an adult. (F-H) Crispant wing phenotypes showing loss of eyespots and pigmentation defects. (I-J) Crispants showing antennal defects (K) Transgenic embryo showing EGFP expression driven by the *Dll319* CRE in mouthparts, antennae, legs, and pleuropodia (red arrows from left to right).

However, one candidate CRE that was open in the distal forewing at scaffold position 150 kb (Fig. 3B) (*Dll319* CRE), returned a unique BLAST hit to the genome. As this region did not contain any repetitive elements, we used CRISPR-Cas9 to disrupt its function. We designed four guide RNAs along its 319 bp length to maximize the likelihood of its disruption (Fig. 3A, Fig. S15). We obtained a variety of different phenotypes that were also observed when targeting exons of the *Dll* gene using CRISPR (*6*): several caterpillars showed a missing or necrotic thoracic leg (Fig. 3C, Fig. S16), adults were missing legs and even a hindwing (Fig. 3D-E), adults lacked eyespots (Fig. 3F-H), adults showed truncated antennae, pigmentation defects, and loss of wing scales (Fig. 3I-J and Fig. S16-S19, Table S1), all having deletions within the CRE of various sizes (Fig. S16). These findings confirm that the *Dll319* CRE is pleiotropic and further suggest that eyespots use the same GRN as antennae in addition to legs and wings.

In order to confirm that the *Dll319* contains a functional and pleiotropic CRE, we cloned a 917 bp region containing this CRE into a *piggyBac*-based reporter construct (*22*) and evaluated its CRE activity in transgenic butterflies. We observed that embryos expressed the reporter gene (EGFP) in antennae, mouthparts, as well as thoracic limbs, indicating that this CRE is sufficient to drive gene expression both in antennae and legs (Fig. 3K and S20). Unfortunately, the loss of this line precluded us from visualizing EGFP expression in eyespots. Using this same cloned region containing the *Dll319* CRE, we also observed pleiotropic CRE activity in antennae, mouthparts, legs, and genitalia, when tested in a cross-species setting with *Drosophila melanogaster* (Fig. S21), suggesting that this region contains an ancestral and pleiotropic CRE present in the ancestors of flies and butterflies.

In order to investigate the extent to which other genes of the eyespot GRN share the same open-chromatin profiles as genes expressed in antennae and in other tissues, we performed an Assay for Transposase-Accessible Chromatin using sequencing (ATAC-seq) with the same tissues used for the transcriptome analysis. A differential accessibility analysis for the open-chromatin regions associated with the eyespot DE genes showed that eyespots shared the greatest number of open-chromatin regions with antennae, as compared to other tissues at the 3-h-pupal stage (Fig. 4G, 4H). The ATAC-seq data also showed that the *Dll319* CRE is open across all different stages and tissues, irrespective of the expression of *Dll* (Fig. 4A), suggesting that pleiotropic CREs may always be open throughout development. To test this, we further targeted a genomic region of *sal (sal740)* that had open-chromatin across most developmental stages using CRISPR-Cas9 (Fig. 4B). We obtained aberrations in caterpillar horns, adult antennae, leg and chevron patterns, as well as missing eyespots and a missing wing (Fig. 4C-4F, Fig. S22), again confirming the presence of a pleiotropic CRE for a gene common to both eyespots and antennae.

**Fig. 4.**
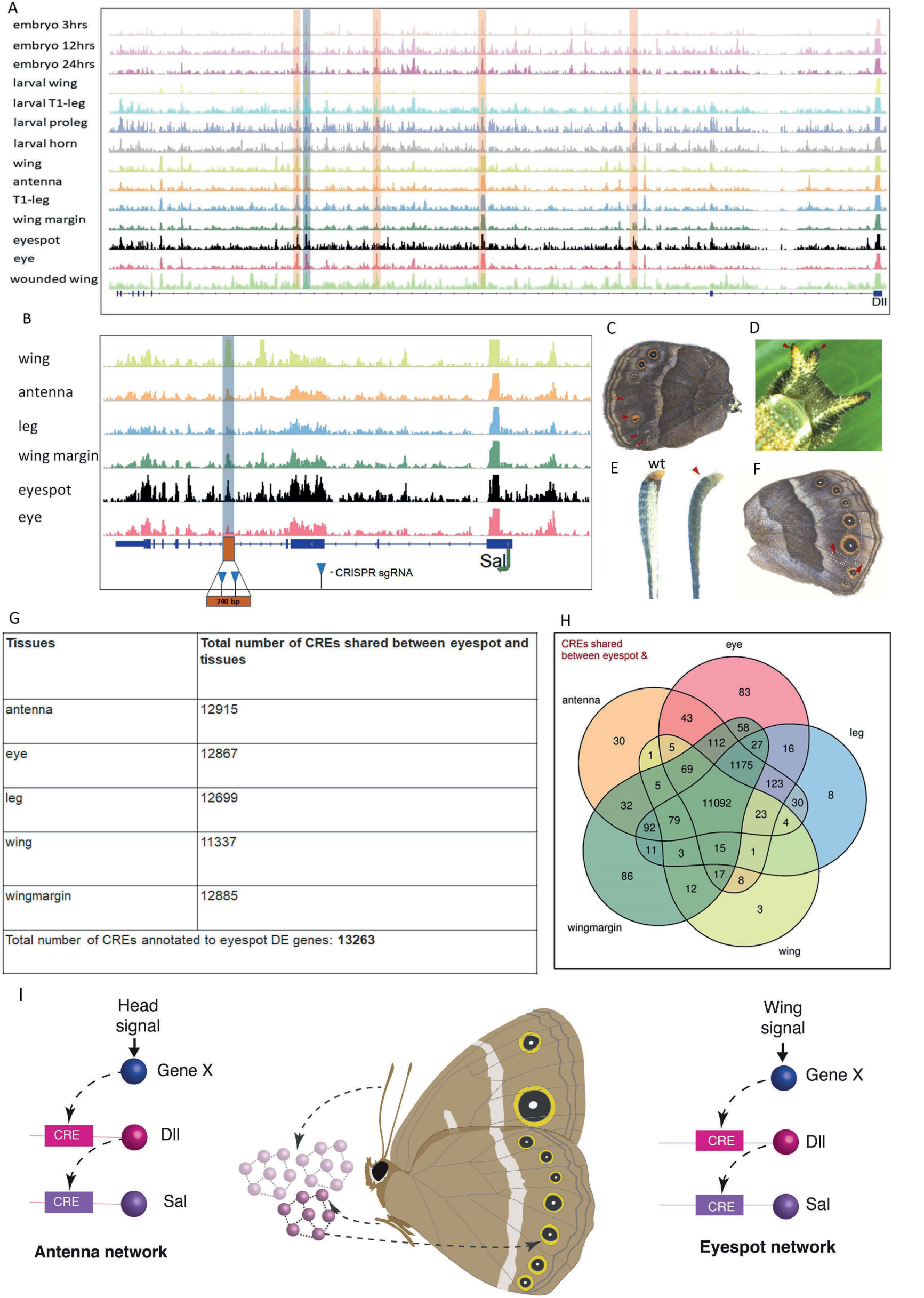
Visualization of open chromatin around *Dll* and *sal* genomic regions for different tissues and identification of a *sal* pleiotropic CRE. (A). ATAC-seq reads around the *Dll* genomic region with highlights in the open regions shared across different tissues (orange) and the targeted *Dll319* (blue). (B). ATAC peak regions from 3hr pupal tissues around the *sal* genomic region with the CRE *(sal740)* targeted highlighted in blue. (C-F). *sal740* crispant phenotypes: Missing and reduced eyespots (C), split horn (D), thinner and discolored antenna compared to wild type (E), lost chevrons in the wing margin and ectopic vein in the Cu2 sector (F). (G). Table with the total number of open peaks associated with eyespot DE genes and number of peaks shared between eyespots and different tissues. (H). Venn diagram showing the number of open chromatin regions shared between different tissue groups. (I). Schematic illustrating the hypothesis that eyespots evolved via co-option of an antennal GRN with genes (*Dll* and *sal*) in the GRN reusing the same CREs in both antennae and eyespot development.

To further confirm that the two CREs *(Dll319 & sal740)* drive *Dll* and *sal* in an endogenous context, we reanalyzed Hi-C data from the wandering larval stage, when Dll and Sal proteins are expressed in eyespot centers (Fig. 2). Using the *Dll319 and sal740* CREs as a bait, we observed that these two sequences physically interact with the *Dll* promotor and *sal* promoter, respectively (Fig. S23).

By exploring the gene expression profile and functional regulatory connections of elements of the eyespot GRN, we showed that eyespots, a morphological novelty in nymphalid butterflies, likely evolved via co-option of the antennal GRN, the oldest urbilaterian appendage. This network, initially deployed in primitive sensory systems, has been subsequently recruited and modified to produce legs (*23*) and perhaps even wings (*24, 25*). We show that the transcriptome profile of eyespots more closely resembles that of antennae compared to any other tested appendage or butterfly tissue. Furthermore, genes known to be critical for eyespot development share the same functional connections as observed in *Drosophila* antennae. Previous studies in *Drosophila* had demonstrated the same CRE driving reporter gene expression in separate traits, and CRE disruptions leading to pleiotropic effects on patterns of CRE activity (*26*). However, here we show, for the first time, that disruptions to two pleiotropic CREs result in the loss of both ancestral and derived traits, which provides uncontroversial evidence for GRN co-option.

The *cis*-regulatory paradigm (*27*) suggests that, when a gene is expressed in a different developmental context, it uses a different CRE for its activation. Here we show that this does not apply to traits that emerge through gene-network co-option, as the recruited network genes are most likely sharing pre-existent regulatory connections (*26, 28*) (Fig. 4I). The origin of novelties has remained an important unanswered question in biology; and here we show that novelties can arise from GRN co-option, which provides a mechanism for complex traits to evolve rapidly from pre-existing traits.

## Acknowledgement

We would like to thank NUS HPC for allowing access to perform the bioinformatic analysis. Tirtha Das Banerjee for his comments on manuscript.

## Authors contribution

SNM, HC, YM, and AM designed and conceived the project. SNM, HC, YM, MDG, MH, GV, SM, KDD performed the experiments. TW and YT critically revised and, SNM, HC, YM, AM analyzed the data and wrote the manuscript.

## Data Availability

All raw Illumina reads of RNA-seq, ATAC-seq and Hi-C are available under NCBI Bioproject (PRJNA685019)

## Materials and Methods

### Butterfly husbandry

*Bicyclus anynana* were maintained in lab populations and reared at 27°C and 60% humidity inside a climate room with 12:12 h light:dark cycle. All larvae were supplied with young corn leaves to complete their development until pupation. Following pupation, the pupae were collected and placed in a separate cage until they emerged. The butterflies were fed every other day with banana on moist cotton in Petri dishes.

### Wing library preparation and FAIRE-seq analysis

Wings were dissected from *B. anynana* at ∼22-26 hours post-pupation. For control input libraries (non-enriched), 2 whole forewings and 2 whole hindwings were pooled. Three FAIRE-enriched libraries were prepared in total, including a forewing distal library (the pupal wing was cut in half and the distal region was used for the library) and 2 hindwing libraries, using both the proximal and distal regions of the wing. All FAIRE-enriched libraries were prepared from 7-8 pooled wing tissues. Libraries were prepared by Genotypic Technology (India) as 75bp pair-end reads and sequenced, using Illumina NextSeq. Raw reads were quality-checked and reads with phred scores >30 were retained for downstream analyses. Following the removal of adapters and low-quality bases, the reads were aligned to a *B. anynana* BAC sequence containing *Dll,* with BWA (0.7.13)(*29*), using the following parameters: –k INT, -w INT, -A INT, -B INT, -O INT, -E INT, -L INT, -U INT. The resulting SAM files were converted to BAM files, using SAMtools-0.1.7a(*30*). The BAM files were converted to sorted BAM, followed by removal of PCR duplicates. The final BAM files were converted to BEDgraph files, using BEDtools-2.14.3(*31*). Peaks were called with MACS2 software(*32*), using the aligned enriched and input (control) files with the q-value (minimum FDR) cutoff to call significant regions. The command bdgcmp script was used on the enriched and input BEDgraph files to generate fold-enrichment and log likelihood scores. This command also removed noise from the enriched sample relative to the control. The BEDgraph files were converted to BigWig files for visualization in Integrative genomic viewer (IGV).

### Identifying *cis*-regulatory elements (CREs) for CRISPR-Cas9 experiments

The FAIRE-seq data were visualized using IGV. All 18 candidate CREs identified around the *Dll* locus were blasted against the *B. anynana* genome in LepBase to verify whether they were unique in the genome. Most of the candidate CREs were not unique and had multiple hits throughout the genome. One of the unique regions, the CRE *Dll319*, was selected as a suitable target for CRISPR knock-out.

### Single guide RNA design and production

Single guide RNA (sgRNA) target sequences for *sal* were selected based on their GC content (around 60%) and the number of mismatch sequences relative to other sequences in the genome (> 3 sites). In addition, we selected target sequences that started with a guanidine for subsequent *in vitro* transcription by T7 RNA polymerase. sgRNA for the *Dll319* CRE were designed using CRISPR Direct(*33*), corresponding to GGN20NGG. We designed 4 guides spanning the length of the CRE (Fig. 3B, Fig. S15, and Table S2). Two guides were designed targeting the *sal740* region (Fig. S24, and Table S2). The sgRNA templates were created by a PCR reaction with overlapping primers, using Q5 polymerase (New England Biolabs). Constructs were transcribed using T7 polymerase and (10X) transcription buffer (New England Biolabs), RNAse inhibitor (Ribolock), NTPs (10 mM) and 600 ng of the PCR template. The final sample volume was 40 μL. Samples were incubated for 16 h at 37°C and then treated with 2 μL of DNAse 1 at 37°C for 15 minutes. Samples were purified by ethanol precipitation, and RNA size and integrity were confirmed by gel electrophoresis.

### *Cas9* mRNA production

The plasmid pT3TS-nCas9n (Addgene) was linearized with XbaⅠ (NEB) and purified by phenol/chloroform purification and ethanol precipitation. pT3TS-nCas9n was a gift from Wenbiao Chen (Addgene plasmid # 46757; http://n2t.net/addgene:46757; RRID:Addgene_46757). *In vitro* transcription of mRNA was performed using the mMESSAGE mMACHINE T3 kit (Ambion). One microgram of linearized plasmid was used as a template, and a poly(A) tail was added to the synthesized mRNA by using the Poly(A) Tailing Kit (Thermo Fisher). The A-tailed RNA was purified by lithium-chloride precipitation and then dissolved in RNase-free water and stored at -80°C. The *Cas9* transcript was used for producing *sal* crispants, and for the analysis of regulatory interactions among *Dll*, *Antp*, and *sal*.

### *In vitro* cleavage assay for the *Dll319* CRE

The sgRNAs were tested using an *in vitro* cleavage assay. Wild-type genomic DNA was amplified using primers that were at least 200 bp from the sgRNA sites. sgRNA (200 ng/μL per guide), Cas9 protein (800 ng/μL) (stored in a buffer containing 300 mM NaCl, 0.1 mM EDTA, 1 mM DTT, 10 mM Tris-HCl, 50% glycerol pH 7.4 at 25°C), NEB buffer 3 (1 μL) and BSA (1 μL) were brought to a final volume of 10 μL with nuclease-free water and incubated at 37°C. After 15 minutes of incubation, the purified amplicon (100 ng) was added to the sample, which was then incubated for an additional 1-2 h at 37°C. The entire reaction volume was analyzed on a 1%-agarose gel. Cas9 protein was purchased from NEB EnGen Cas9 NLS. The cleavage assay confirmed that each guide successfully cleaved the PCR amplicon.

### Embryo injections

Wild-type lab populations of *B. anynana* adults were provided with corn plants to lay eggs. The eggs were collected within 1.5 h of oviposition and placed onto 1-mm-wide strips of double-sided tape in plastic Petri dishes (90 mm). Cas9 protein (final concentration 800 ng/μL) and sgRNA (final concentration 200 ng/μL per guide) for all 4 guides were prepared in a total volume of 10 µL and incubated for 15 min at 37°C prior to injection along with 0.5 µL of food dye to improve visualization of the injected sample into the embryos. For *sal* crispants, Cas9 mRNA (500 μg/μL final concentration) and sgRNA (500 μg/μL final concentration) were injected along with one tenth of the volume of food dye. For *sal740* CRE, eggs were injected with the mix of Cas9 protein (final concentration 800 ng/μL) and sgRNA (final concentration 400 ng/μL per guide). The injection mixture was kept on ice after the incubation and prior to injection. Embryo injections were carried out by nitrogen-driven injections through glass capillary needles. Injected eggs were stored in closed Petri dishes containing cotton balls that were dampened daily to maintain humidity. The hatched larvae were reared in small paper cups for 1 week and then moved to corn plants to complete their development. Tables S1, S3 and S4 summarize the injection results.

### *In vivo* cleavage assay and genotyping of *sal* crispants

Genomic DNA was extracted with a SDS-based method from a pool of 5 injected embryos that did not hatch. About 250 bp of sequence spanning the target sequence was amplified with PCRBIO Taq Mix Red (PCRBIOSYSTEMS), and PCR conditions were optimized until there were no smears, primer dimers, or extra bands. Primers are listed in Table S2. The PCR products were purified with the Gene JET PCR purification kit (Thermo Fisher). Two hundred nanograms of PCR product were denatured and re-annealed in 10x NEB2 buffer. One microliter of T7 endonuclease Ⅰ (NEB) was added to the sample, while 1 µL of MQ water was added to a negative control. Immediately after the incubation for 15 min at 37°C, all the reactions were analyzed on a 3% agarose gel. Amplicons that showed positive cleavage from the T7 endonuclease Ⅰ assay were subcloned into the pGEM-Teasy vector (Promega) through TA cloning. For each target, we picked 8 colonies, extracted the plasmid with a traditional alkali-SDS method, and performed a Polyethylene glycol (PEG) precipitation. Sequence analysis was performed with the BIGDYE terminator kit and a 3730xl DNA Analyzer (Thermo Fisher).

### Screening and genotyping *Dll319* crispants

Newly emerged caterpillars were screened under a microscope to look for developmental defects affecting any regions where *Dll* is expressed, such as the thoracic legs, mouthparts, and prolegs. Any caterpillars exhibiting defects were imaged and reared individually in paper cups until the butterflies eclosed. Caterpillars that died were immediately frozen for DNA isolation and genotyping. All other surviving caterpillars with no apparent developmental abnormalities were reared in groups on corn plants and fed *ad-libitum* every 2 days until pupation. The eclosed butterflies were frozen individually at -20°C. Each butterfly was carefully screened under a microscope and examined for asymmetric crispant phenotypes, focusing particularly on phenotypes expected for a *Dll* knock-out, such as appendage, eyespot, or pigmentation defects.

### Colony PCR to identify CRE deletions

For selected crispants, genomic DNA was extracted from the thorax (E.Z.N.A tissue DNA kit) and used for PCR to prepare samples for genotyping. The samples were visualized on a gel to confirm the correct size band and the PCR product was purified using a Thermo Scientific PCR purification kit. The DNA was cloned into a pGEM T-Easy Vector (Promega) and the plasmid was transformed into DH5 alpha *E. coli.* White colony selection was used for colony PCR. The bands were visualized on a 1%-agarose gel to look for bands with shifts relative to the WT band. PCR products from colonies showing evidence of a deletion were submitted for Sanger sequencing PCR (Axil Scientific, Singapore), including a sample that was amplified from *B. anynana* wild-type genomic DNA.

### Butterfly enhancer reporter assay

A 917 bp region containing the *Dll319* CRE was cloned into the piggyGUE vector via the GATEWAY technology (Thermo Fisher). piggyGUE is the EGFP version of piggyGUM, the piggyBac-based reporter construct that was previously published(*22*). The details of piggyGUE will be published elsewhere (Deem and Tomoyasu, unpublished). The 917 bp region was amplified from *B. anynana* wild-type genomic DNA using a primer containing CACC at the 5’ end for directional cloning. The PCR product was cloned into the pENTR vector and further cloned into the piggyGUE vector via a LR reaction, as described by Lai et al., 2018(*22*). Four microliters of the LR reaction mix were used for bacterial transformation. After sequence analysis to confirm the presence of SNPs in the *Dll319* CRE, plasmid DNA was amplified, using a Midiprep kit (Qiagen). The piggyGUE *Dll319* CRE plasmid was diluted to 1µg/µL and mixed in a 1:1 ratio with a hyperactive *piggyBac* transposase plasmid(*34*). Embryos (n=550) were collected from *B. anynana* butterflies reared at 27°C and were injected ∼1-hour after egg laying with the plasmid solution and a small amount of food dye, using a glass injection needle and nitrogen gas pressure. Eggs were transferred in a Petri dish to a chamber and kept moist to prevent dehydration. From this batch of eggs, 40 caterpillars hatched and were reared in paper cups during the first week and then transferred to cages with corn plants to complete their development. At all stages, caterpillars were fed corn *ad-libitum*. From this batch of caterpillars, 19 reached adulthood (10 females and 9 males). These butterflies were evenly distributed into 4 cages (∼5/cage) and placed with respective wild-type males and females for breeding. We were unable to observe any *dsRed* signal (the positive marker of transgenesis driven by the 3xP3 promoter) in the eyes of the caterpillars from the F1 or F2 generation, despite ubiquitous *dsRed* signal in some 1^st^-intar larvae (only) of the F1 generation, which were used later for outcrossing to wild-type individuals. This ubiquitous signal was not observed again in the offspring of these larvae. We collected eggs from the F3 generation and dissected some embryos for EGFP antibody staining. Two out of the four dissected embryos did show expression of EGFP driven by the *Dll319* CRE in the embryonic antennae, mouthparts, thoracic legs and pleuropodia (Fig. 3K, Fig. S10). Subsequent hemolymph PCR genetic screening in individuals of the 4^th^ generation failed to identify additional positive individuals and the line was lost.

### *Drosophila* enhancer reporter assay

The same 917 bp sequence that contained The *Dll319* CRE was directionally cloned into pENTR-D, then GATEWAY cloned into the piggyPhiGUGd, the Gal4-delta version of the previously reported piggyBac-based reporter construct(*22*). piggyPhiGUGd also has an attB site, allowing phiC31 transgenesis. The detail of piggyPhiGUE will be published elsewhere (Deem and Tomoyasu, unpublished). For *Drosophila* transgenesis, the piggyPhiGUGd *Dll319* CRE construct was transformed into the attP2 site (68A4) through phiC31 integrase-mediated transgenesis system with EGFP as a visible marker (BestGene *Drosophila* transgenic service). Established transgenic flies were crossed with G-TRACE(*35*) to visualize the tissues with CRE activities.

### Antibody staining of *B. anynana* embryos and wings

Two-day-old embryos, as well as fifth-instar larval and pupal wing tissues were dissected in PBS buffer under the microscope. The samples were fixed in 4% formaldehyde/Fix buffer (0.1 M PIPES pH 6.9, 1 mM EGTA pH 6.9, 1.0% Triton x-100, 2 mM MgSO4) for 30 min on ice. The samples were washed with 0.02% PBSTx (PBS + Triton x-100) 3 times every 10 min, and then blocked in 5% BSA/PBSTx for 1 h. The samples were then incubated in 5% BSA/PBSTx with the primary antibody, and incubated at 4°C overnight. As primary antibodies, we used a rabbit polyclonal anti-Dll antibody (at 1:200, a gift from Grace Boekhoff-Falk), a mouse monoclonal anti-Antp 4C3 antibody (at 1:200; Developmental Studies Hybridoma Bank), a rabbit anti-Sal antibody (at 1:20,000 for wings and pupal tissues, and 1:2,000 for embryos; de Celis et al., 1999), and a rabbit anti-EGFP antibody (at 1:200; Abcam ab290) for the transgenic embryos at 24h (n=4) and wt controls. For double staining, we added two primary antibodies to the same tube. The wings were washed with PBSTx 3 times every 10 min. Then, we replaced the PBSTx with 5% BSA/PBSTx to block for 1 hour, followed by the incubation with the secondary antibody (1:200) in 5% BSA/PBSTx at 4°C for 2 h. The wings were washed with PBSTx 3 times every 10 min, followed by mounting the wings in ProLong Gold Antifade Mountant (Thermo Fisher). The images were taken under an Olympus FV3000 Confocal Laser Scanning Microscope.

### Sample collection and library preparation for RNA sequencing

In order to identify gene expression patterns specific to eyespot formation on the developing wings, we extracted RNA from sixteen different tissue types at four developmental time points: 3-4-hour-old, 12-13-hour-old, and 24-25-hour-old embryos; T1 legs, prolegs, forewings, and horns from wandering caterpillars; T1 legs, antennae, forewings, forewing margins, eyes, eyespots, and two control tissues adjacent to eyespot centers from 3-h-old pupae (**Fig. 1A**). For wing wounding experiments, we poked one wing between 17 to 18 h after pupation in two different places in the M3 sector, using a fine tungsten needle with a diameter of 0.25 mm and 0.001 mm at the tip (FST- 10130-10). We collected the wings 6 hours later, which corresponds to 23-24 h after pupation (Monteiro et al 2006). We performed the experiments with four biological replicates for each tissue type with 10 to 25 female individuals in each replicate (both left and right tissues were used, except for the wounded pupal wings, where a single wing was used) (Table S5). Total RNA was extracted in 70 µL of nuclease-free water, using Qiagen RNA Plus Mini Kit. RNA quantity and integrity were measured using a Nanodrop and an RNA Bleach gel (Aranda et al 2013). RNA libraries were prepared, using the Truseq stranded mRNA kit from Illumina. Forty million reads were sequenced for each replicate, using Novoseq 6000 with 150bp paired-end and an average insert size of 250-300 bp. Library preparation and sequencing were carried out at AIT Novogene, Singapore. In order to avoid batch effects, we randomized the sample extraction and RNA isolation, such that two replicates of the same group were never extracted at same time.

### RNA-seq analysis

The raw RNA-seq data were quality-controlled and filtered. Adapter sequences and reads with low quality (less than Q30) were trimmed, using bbduk scripts (ktrim=r, k=23, mink=11, hdist=1, tpe, tbo, qtrim=rl, trimq=30, minlen=40). In order to remove any bacterial contamination in the samples, we used the bbsplit script, which is a part of the bbmap tools (*36*). All bacterial genomes were downloaded from NCBI (last downloaded in June 2018), and the reads were mapped to the bacterial genomes, using bbmap. Only reads whose pairs also passed through the filter were further analyzed. To remove any ribosomal RNA sequences from the RNA-seq data, the reads were aligned to the eukaryotic rRNA database available in sortmeRNA (*37*). The processed reads from different samples were then mapped to the BaGv2 genome, using hisat2 (*38*) (mapping statistics in Table S6), resulting in bam files that were sorted by genomic positions, using samtools (*30*). They were used as inputs in StringTie (*38*) to create the initial transcriptome assembly with 71,042 transcripts, which was used to annotate the genome using Maker v.3 (*39*), resulting in 18,196 genes with 29,389 transcripts.

### RNA-seq differentially expressed (DE) gene analysis

A read count matrix of the annotated genes was obtained for the samples using StringTie (*38*). We used the GO terms to filter out any ribosomal genes before obtaining the read counts. This approach led to the removal of 496 genes to a final set of 17,700 genes, which was used throughout the analysis. Correlations between the replicate samples was analyzed using DESeq2 (*8*) with a sample distance matrix. One of the antennal samples was removed due to its poor correlation with its other biological replicates. The remaining samples were used for the downstream analyses (Fig. S25).

### Identifying eyespot-specific DE genes

To identify eyespot-specific genes, a pairwise DE analysis was performed between eyespot and control adjacent tissues, Nes1 and Nes2, using DESeq2 (Fig. 1A, Fig. S2). Common genes upregulated and downregulated between eyespot vs. Nes1 and eyespot vs. Nes2 with an adjusted P-value (padj) of 0.05 were chosen as eyespot-specific DE genes (Spreadsheet S1).

### RNA hierarchical sample clustering

In order to identify the tissue with the closest gene expression profile to eyespot, we used all tissue samples except the eyespot control tissue samples. DE analysis between the multiple tissues was performed, using **run_DE_analysis.pl** script provided in Trinity tool, using DESeq2 as the method of choice for this analysis (*40*). Hierarchical clustering was performed for the different tissues, using genes that showed a log2fold change of |2| and padj value of 0.001, as in Fisher et al (2020) (*41*). Clustering was performed using an Euclidean distance matrix derived, using the DE genes for the tissues with the hclust function in R(*42*). The pvclust package(*9*) in R was used to calculate the uncertainty in the hierarchical clustering with a 1000 bootstrap value.

### ATAC-seq library preparation

We prepared ATAC libraries for the same set of tissues as we did for the RNA-seq experiment, except for the eyespot control tissues (Table S7). Library preparation failed for a few groups leading to 2 to 4 biological replicates per group. Tissues were collected, flash-frozen in liquid nitrogen and stored in -80֯ C, before we extracted nuclei and prepared the libraries. We used 10 to 25 individuals and approximately 80,000 nuclei per replicate. Libraries were prepared as described in the Omni-ATAC protocol (*43*) with slight modifications. Individual tissues extracted at different time periods during the process were randomized and pooled into each replicate before extracting the nuclei. The tissues were thawed and homogenized in 2 mL of ice cold 1X homogenization buffer (HB) in a 2-mL-glass douncer. Homogenization was performed by 10 strokes with pestle A, followed by 15 strokes with pestle B. The homogenized mixture was left on ice for 2 min before filtering it through a 100-µm- nylon mesh filter into a DNA “low bind” 2-mL Eppendorf tube (Z666556-250EA). The filtered mixture was centrifuged at 2500 rpm, and the pellet (the nuclei) was collected along with 50 µL of the solution at the bottom, keeping unwanted cytoplasmic RNAs in the top layers. The filtered nuclei were diluted in ATAC-Resuspension Buffer (RSB buffer), and 10 µL of the solution were used to count the nuclei, using a hemocytometer. Approximately 80,000 cells were used for each replicate to prepare the libraries. The tagmentation enzyme (TDE1) was obtained from Illumina (Illumina tagment dna tde1 enzyme and buffer smaller kits – 20034197). As the concentration of the TDE1 and cell number greatly affect the identification of open chromatin regions, we estimated the amount of enzyme needed for each reaction, using the formula: volume of enzyme = genome size of *B anynana* [475MB] * number of cells [80,000] *2.5/ (genome size of humans [3200MB] *50,000). We used 0.65 µL (final concentration of 10.4 nM) of enzyme for each reaction. The Omni-ATAC transposition reaction was carried out as follows: 80,000 cells suspended in ATAC-RSB buffer were centrifuged at 2500 rpm for 10 min at 4° C. The supernatant was removed, and the nuclei-containing pellet was kept. To perform the cell lysis, 50 µL of ice-cold ATAC-RSB were added to the pellet, along with 0.1% NP40, 0.1% Tween 20, and 0.01% Digitonin. The mixture was incubated for 3 min on ice. Subsequently, 1 mL of ATAC-RSB buffer containing only 0.1% Tween and no NP40 nor digitonin was added, and the mixture was centrifuged at 2500 rpm. The supernatant was discarded, and the pellet was retained, to which 50 µL of transposition mixture (6.5 µL 2x TD buffer, 0.65 µL transposase (10.4 nM final concentration), 16.5 µL PBS, 0.5 µL 1% digitonin, 0.5 µL 10% Tween-20, 25.35 µL H2O) were added. The reaction was incubated for 25 minutes at 37° C at 1000 rpm in a thermomixer. After the transpositions and tagmentation occurred, the samples were prepared for sequencing by adding Illumina/Nextera adapters with dual indexing and further PCR amplified for 10 cycles. The PCR products were purified, using a Zymo-DNA Clean & Concentrator-5 kit, and the DNA fragments were size-selected between 50 – 1500 bp, using the ProNex Size-Selective Purification System (NG2002) from Promega. The samples were sequenced, using Novoseq 6000 with an average read depth of 30 million and 2x50 bp paired end reads by AIT Novogene, Singapore.

### ATAC-seq peak calling

ATAC reads were processed, using bbduk scripts from bbmap tools, to remove any adapters. The reads were mapped to the BaGv2 genome, using bowtie with the -x 1500 and -m1 parameters. Only reads with insert sizes of 1500 bp or less and only those mapping to a unique region of the genome were mapped. Reads mapped to the mitochondrial genome were removed, using samtools idxstats, and reads marked for PCR duplicates were also removed, using GATK Markduplicates. We kept only paired-end mapped reads with a phred quality score of Q20 and above for downstream analysis. Because the Tn5 transposase binds to DNA as a dimer and inserts adapters of 9 bp in length at the insertion sites, the start sites of the mapped reads were adjusted to an offset of +4 bp in the forward strand and -5 bp in the reverse strand. The bam files were converted to bed files, using Bedtools (*31*), and we used F-Seq (*44*) to call peaks for each sample. Bedtools intersect was used to identify the common set of peaks for each tissue type. Peaks from all samples were merged if they were separated by 50 bp, using Bedtools merge to create 313,425 consensus peaks used for the downstream analyses. Featurecount from the Subread package(*45*) was used to extract a read count matrix corresponding to the consensus peaks for all samples. The FRiP score, which is defined as the fraction of all reads that are mapped to peaks across the entire genome, was used to measure the quality of the ATAC-seq data. Our ATAC-seq data showed a median FRiP score of 0.846, which is higher than the ENCODE standard (>0.3) for the fraction of reads falling into peaks (Table S8). And deepTools (*46*) was used to access the sample correlation between the replicates and quality of the libraries (Fig. S26).

### ATAC-seq differential peak analysis

Differential peak analysis was performed using DESeq2. Peaks were considered differentially accessible with a padj value of 0.05. We also mapped the *Dll319* peak identified from the FAIRE data to the BaGv2 genome, using blastn, to identify its position in the new genome assembly and test whether the ATAC-seq analysis was also able to identify it. To identify potential CREs for *sal,* ATAC peaks from 3hr pupal tissues were visualized using IGV and we targeted one potential candidate region (sal740) within the intronic region of *sal* gene loci which is open across almost of the tissues.

### Hi-C analysis and Virtual 4C

The Hi-C library used for scaffolding the *B. anynana* genome was reanalyzed, using the *Dll319* and sal740 region as bait, to verify whether these regions interacted with the promoter of *Dll and sal* respectively. Libraries were mapped to the BaGv2 assembly, using Juicer (*47*). We used the contact map obtained from Juicer to construct a virtual 4C plot for the window around the *Dll319* and sal740 regions by placing reads in a 3-kb bin, using the script from Ray et al., 2019 (*48*).

### Screening and genotyping *sal740* crispants

Caterpillars that emerged were carefully screened under the microscope for any defects in their body, especially in the head region where *sal* expression is observed. Individuals showing any abnormalities were imaged and grown separately in a cup whereas all others were grown in a separate cage. Adults were immediately frozen at -20֯ C after emergence and screened later under a microscope for any defects in eyespots, wings, legs and in antenna.

### Hi-C Genome Assembly

Eggs were collected from a single pair of mated *B. anynana* butterflies and reared. Eighteen female siblings were harvested at the wandering stage for DNA extraction. Guts were removed, and the samples were immediately flash-frozen in liquid nitrogen and stored at –80 °C before the samples were sent to Dovetail Genomics to perform Chicago and Hi-C library preparation and analysis. The Chicago library preparation uses *in vitro* chromatin fixation, digestion, and crosslinking of regions in the genome that are close to each other in terms of 3D chromatin architecture. In order to sort and scaffold the genome, 233 million reads (2x150bp) were sequenced from the Chicago library and mapped to the previously published *B. anynana* genome (v1.2) with 10,800 scaffolds(*49*). The HiRise pipeline was used to identify mis- assemblies, to break the scaffolds, and to sort the scaffolds. Only scaffolds greater than 1kb in length (n=5027) were used because scaffolds needed to be long enough for the read pairs to align and be scaffolded in accordance with the likelihood model used by HiRise. Next, 153 million reads (2*150) sequenced from the Hi-C library were mapped to the genome assembly output generated from the Chicago-HiRise pipeline to identify any mis-assemblies from the Chicago pipeline and correct them to produce a final genome assembly of high contiguity.

The genome assembly obtained from the HiC pipeline was ordered, using the available linkage information from Beldade et al., (2009)(*50*), using Chromonomer(*51*). Two hundred eighty-nine SNP FASTA sequences were mapped to the Hi-C assembly, using blastn to identify their corresponding positions in the Hi-C genome. Using the SNP position obtained from blastn, a list describing the genetic map was manually created, which later passed through Chromonomer to sort the Hi-C assembly resulting in the final assembly (**BaGv2**) that was used for the current study. The BUSCO score(*52*) was used to check for the completeness of the gene sets in the assembly.

#### Genome Annotation

The genome was repeat-masked for transposable elements, small repeats, and tandem repeats before annotation as described in Nowell et al., 2017 (*49*) The soft repeat-masked genome was annotated, using four rounds of Maker v.3 (*39*). The transcriptome assembled from the RNA-Seq data and gene sequences annotated from the previous version of the genome were combined and used as transcripts for the species, with transcriptome and protein sequences from *Pieris rapi, Junonia coenia*, and *Bombyx mori* as relative transcripts and protein homology evidences for the first round of gene predictions. Output gene predictions from each round were used as input for the next round. Snap and Augustus were used for the second round of gene predictions, followed by Genemark for the third round of gene modelling. Then we performed one final round of Snap and Augustus predictions. The minimum length of 35 amino acids was set for gene predictions. The predicted gene models were kept for genes that had an Annotation Edit Distance (AED) score of < 1 and/or had a gene ontology obtained from Interproscan (*53*). This resulted in 18,189 genes with 29,490 transcripts. In order to correct the annotations and produce a standardized gff3 file, the gff file obtained from Maker was run through agat_convert_sp_gxf2gxf.pl script, which is a part of AGAT tools (*54*). This step resulted in the removal of 82 identical isoforms and added the missing gene features, leading to a total of 18,196 genes with 29,389 transcripts. Functional annotation was performed by locally blasting the transcripts against a non-redundant (nr) protein database, using diamond blast (*55*), and a gene ontology analysis was performed using Interproscan in Blast2Go(*56*). Finally, the blast results were merged with the interproscan results in Blast2Go to produce a final functional annotation for the genome.

## Supplementary Results

### *Bicyclus anynana* Hi-C Genome Assembly

The published version of the *B. anynana* genome assembly (475 MB) contains 10,800 scaffolds with an N90 value of 99.3 kB (*49*). To improve the current assembly, we performed scaffolding, using a two-step approach, one with Chicago-HiRise followed by Hi-C- based scaffolding. Chicago-HiRise scaffolding performed on the published version resulted in 3512 new joins with 634 breaks remaining in the genome, raising the N90 value to 840 kB. The Hi-C scaffolding that followed corrected mis-joints from the Chicago-HiRise scaffolding by creating four new breaks but made 512 new joints, improving the N90 value further to 12.073 MB and placing 98% of the bases (467.62 MB) into 28 scaffolds, achieving a near-chromosomal level assembly. Following the Hi-C assembly and using the linkage map from Beldade et al. (2009)(*50*) obtained for the 28 chromosomes of *B.* anynana, we produced one manual break and one joint to achieve congruence of the two data sets. We were able to map 171 markers out of the 289 markers from the 28 linkage groups (*50*) in the genome, resulting in an ordered chromosomal level assembly of 475.8 MB

## Supplementary Figures

**Fig. S1.**
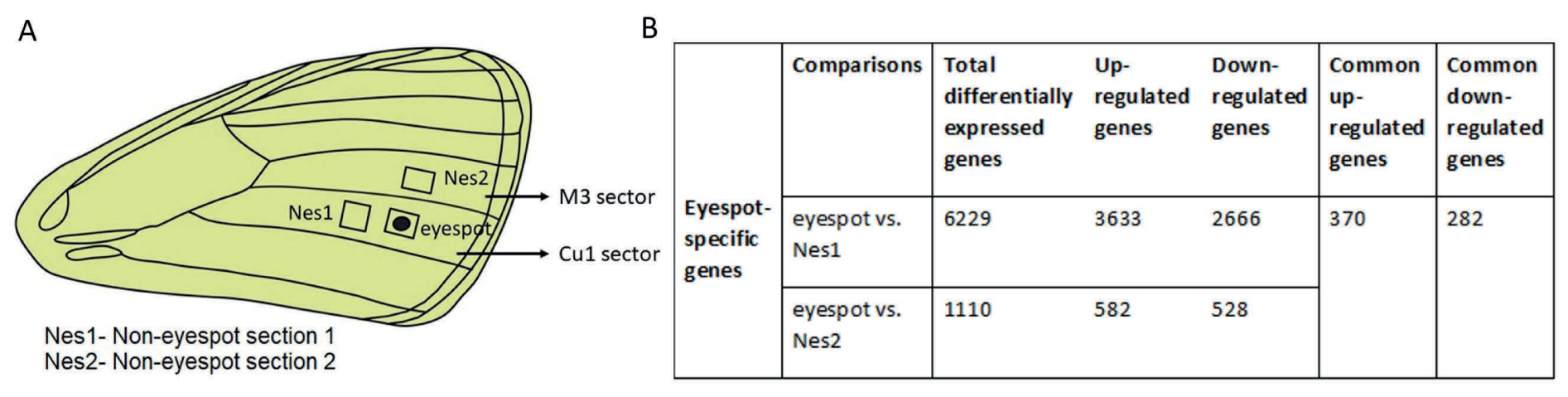
Tissue selection to identify eyespot-specific DE genes. (A) Forewing image highlighting the regions chosen to perform DE analysis to identify eyespot genes. Nes1 and Nes2 are two control tissues representing tissue from the same and from a more anterior wing sector, where no eyespots develop. (B) Table Showing the number of genes differentially expressed (DE) in each of the comparisons, leading to 652 eyespot-specific DE genes that were in common across both comparisons.

**Fig. S2.**
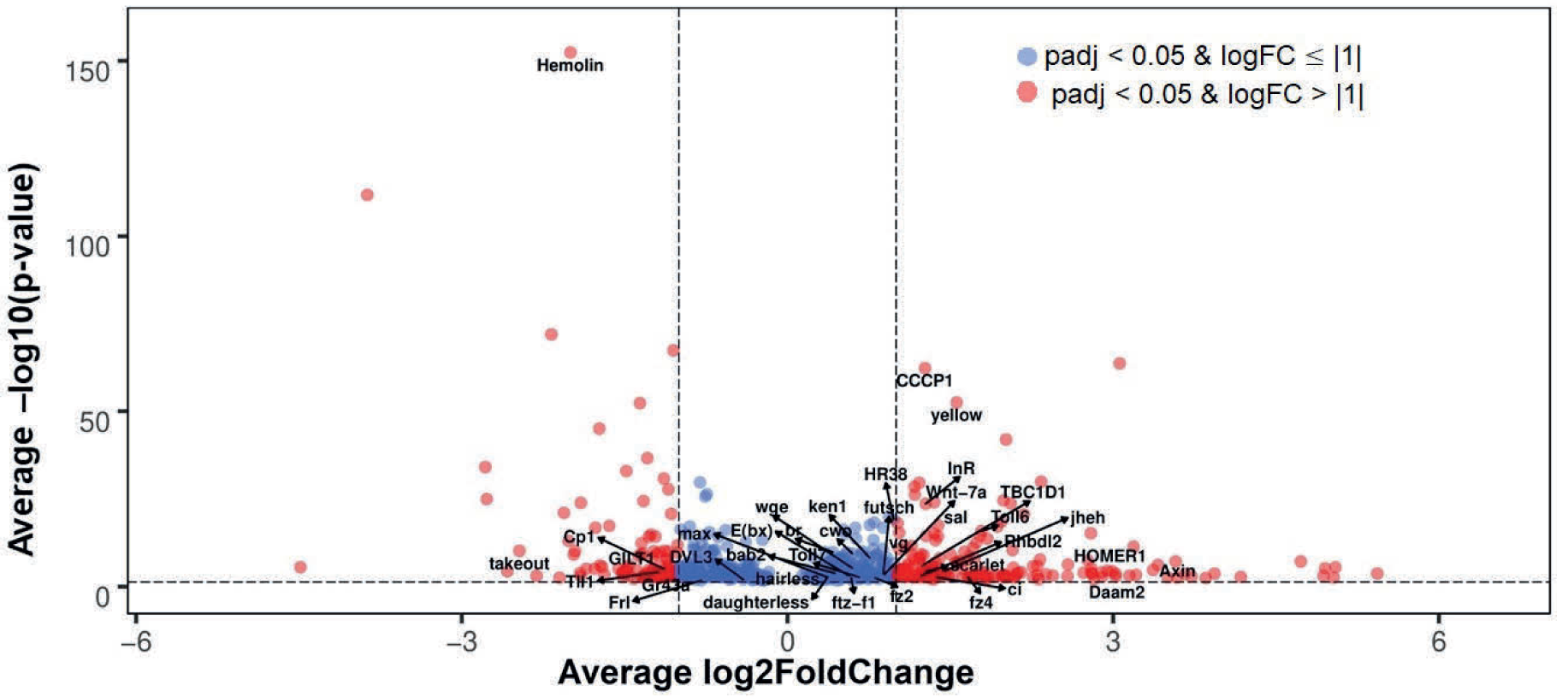
Set of DE genes between the “eyespot” and “control” wing tissues (with adjusted *p*- values <0.05). Up-regulated genes show a positive X-axis value, while down-regulated genes show a negative X-axis value. The log2FC and *p*-values were averaged among the two control tissue comparisons.

**Fig. S3.**
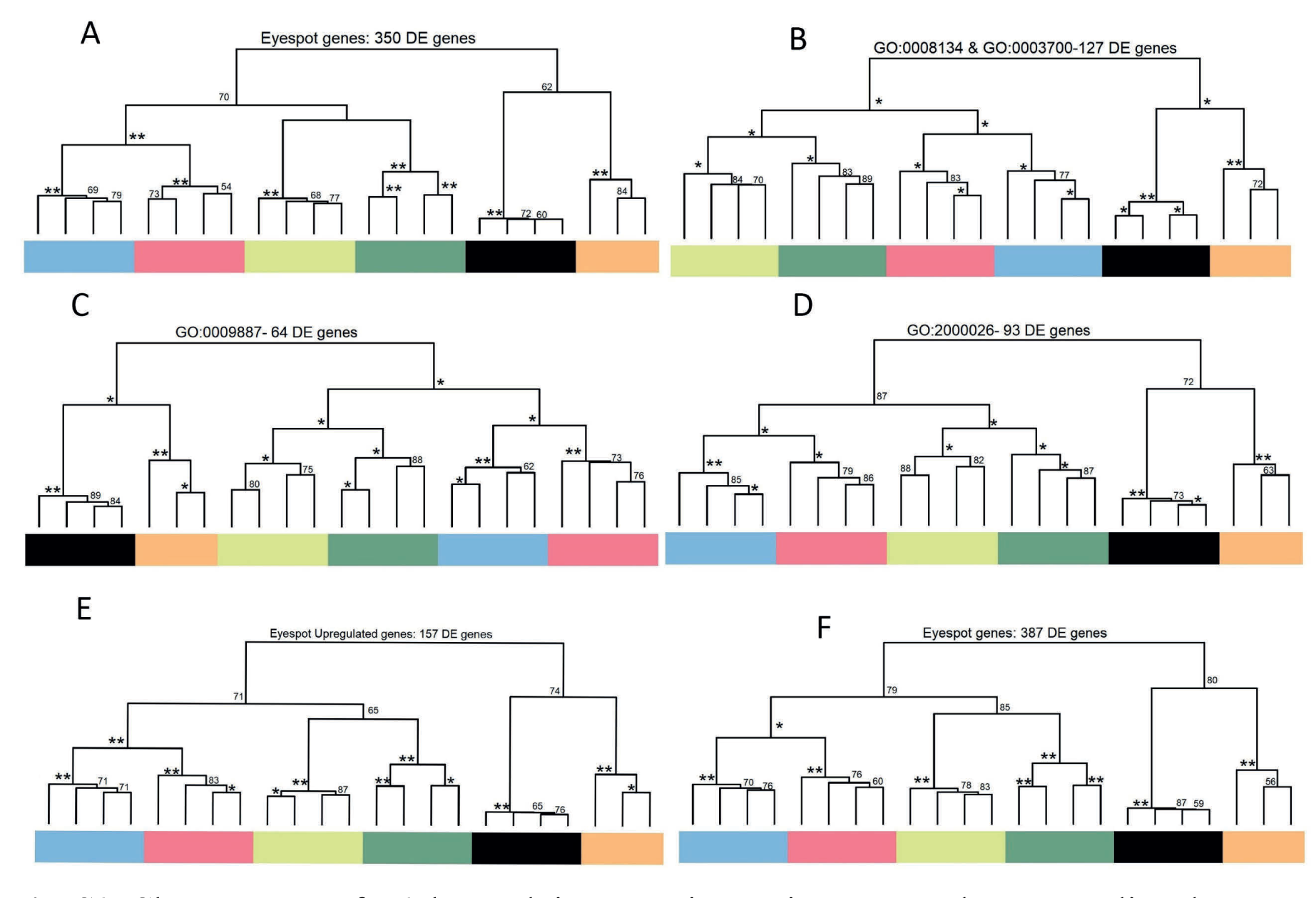
Character trees for 3-h pupal tissues, using various gene subsets revealing that eyespot gene expression patterns cluster with those of antennae. “DE genes” represents the number differentially expressed genes (padj < 0.001 and log2FC ≥|2|) from the initial subset of genes. Trees constructed using: A. eyespot-specific genes (652 genes), B. Transcription factors and co-factors (336 genes), C. genes enriched for GO terms associated with animal organ morphogenesis (108 genes), D. genes enriched for GO terms corresponding to anatomical structure formation involved in morphogenesis (165 genes), E. genes up-regulated in eyespots (370 genes), F. combined eyespot-specific DE genes predicted in this study as well as those published in Ozsu and Monteiro 2017 (*10*) (a total of 775 genes). In all six trees, eyespots gene expression patterns cluster with those of antennae and form an outgroup in the tree with another clade where eyes cluster with legs, and wings with wing margins.

**Fig. S4.**
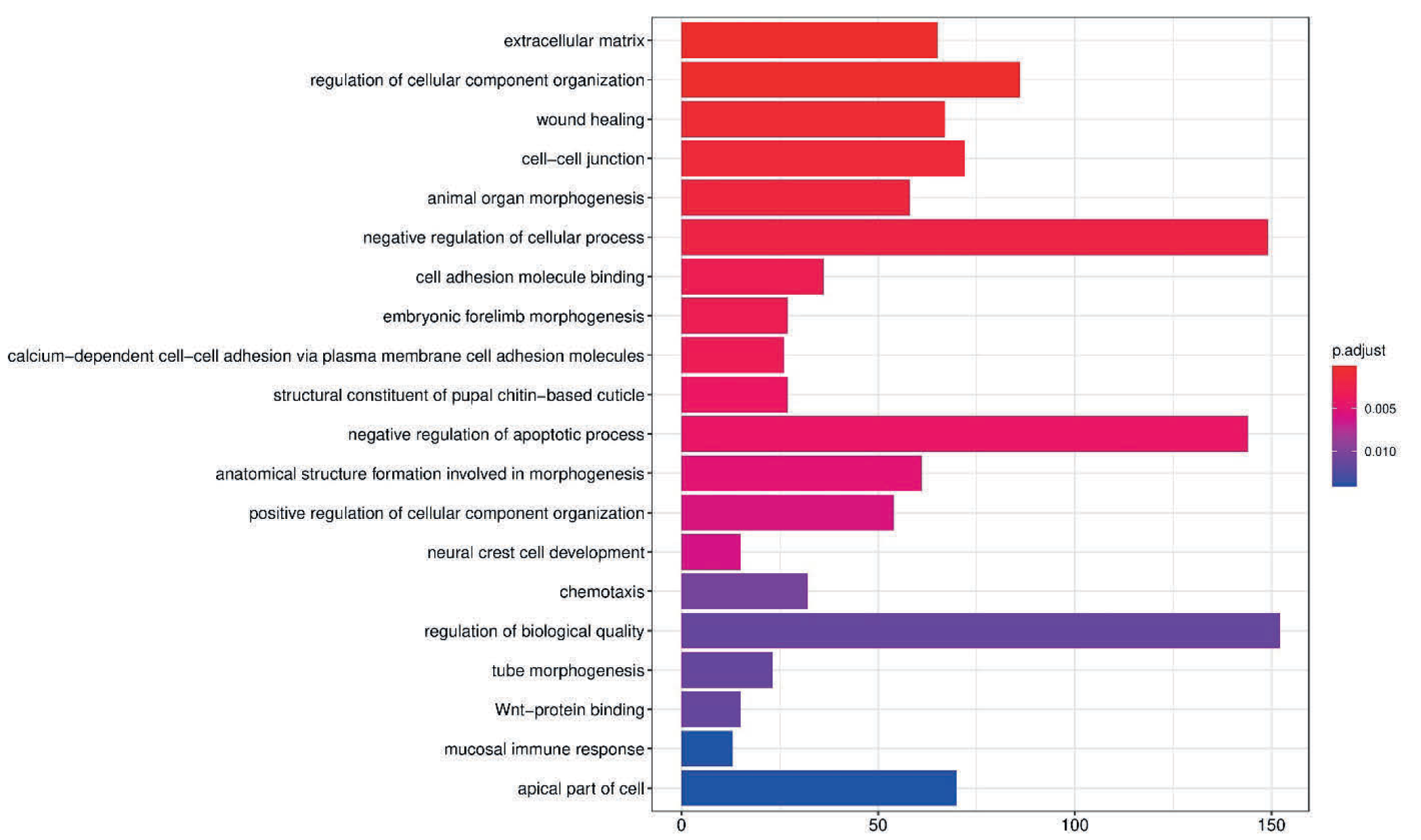
Gene-set-enrichment plot for the 3839 DE genes from 3-h pupal tissues. The gene-enrichment analysis shows the various functions DE genes are involved.

**Fig. S5.**
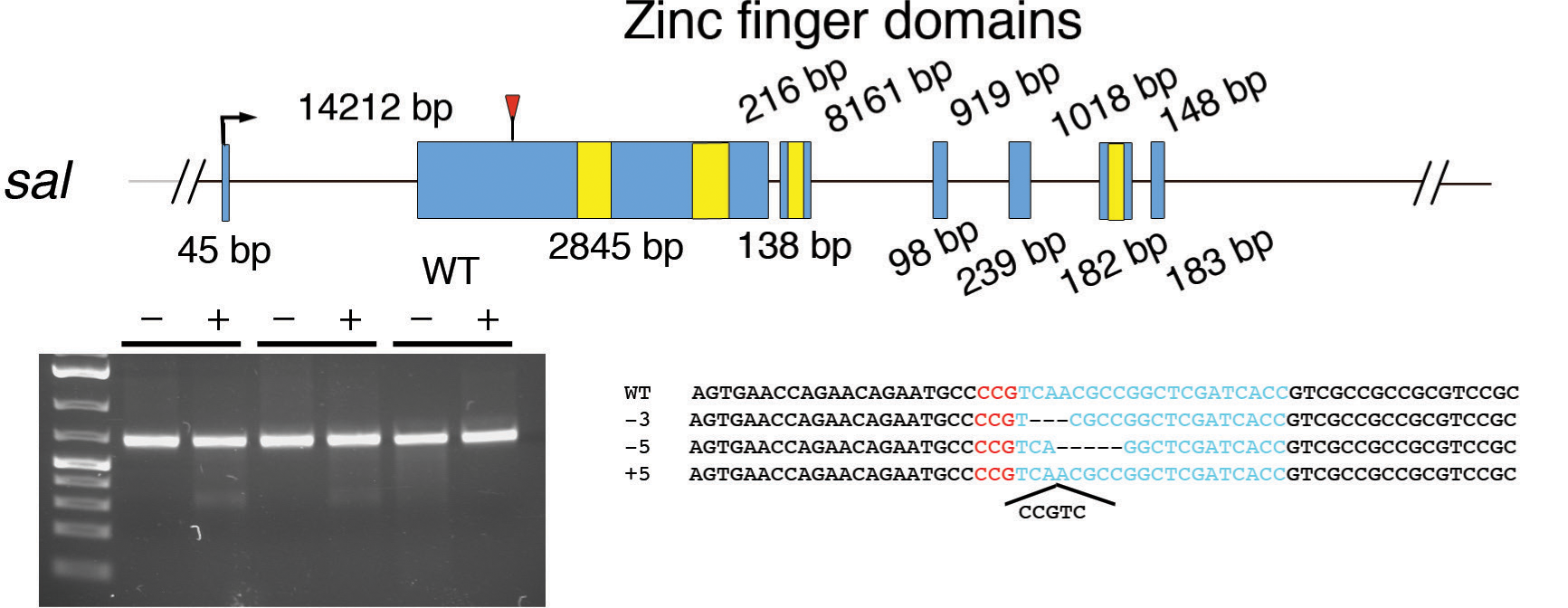
T7 endonuclease I assay and sequence analysis for CRISPR-Cas9 mutations in *sal*. Schematic representation of the *sal* gene structure. Blue boxes indicate exons, and yellow-coloured regions inside exons indicate functional domains. Each functional domain was annotated using a conserved-domain search at NCBI. Red arrowheads indicate the CRISPR target region. The gel shows the result of a T7 endonuclease assay performed on embryos after injection of sgRNA and Cas9 mRNA or protein or performed on Wt embryos (last two lanes). We performed the assay with two different samples for technical replicates. “Minus” lanes indicate a negative control, where T7 endonuclease was not added to the reaction. “Plus” lanes indicate the presence of T7 endonuclease. The expected sizes of digested bands were observed only from the lanes containing the T7 enzyme. Sanger sequence results indicate that an indel mutation was generated around the target site. Blue-coloured sequences indicate the sgRNA target sequence, and red-coloured sequences indicate the PAM sequence.

**Fig. S6.**
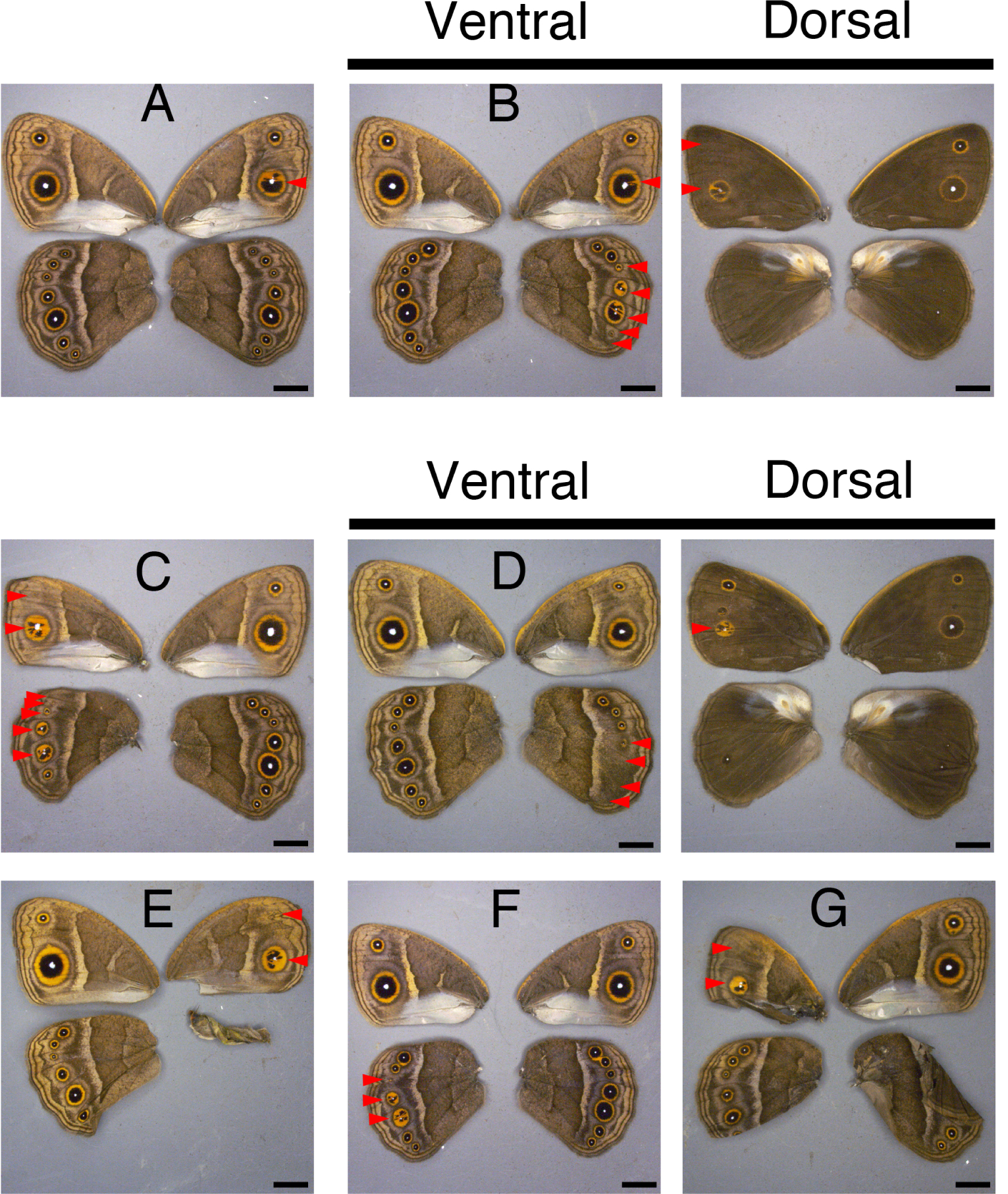
*sal* crispant phenotypes. (A) The Cu1 eyespot on the right forewing showed a transformation of orange scales into black scales. (B) The Cu1 eyespot on the right forewing and the M1, M2, M3, and Cu1 eyespots on the right hindwing showed transformation of black into orange scales. The Cu2 eyespot got reduced in size, and the A1 eyespot on the right hindwing disappeared. On the dorsal side, the M1 eyespot on the right forewing disappeared, but the Cu1 eyespot on the right forewing showed a transformation of black into orange scales. (C) The M1 eyespot on the left forewing was reduced in size, and the Cu1 eyespot showed a transformation of black into orange scales. On the hindwing, the Rs eyespot disappeared, and the M1, M2, M3, and Cu1 eyespots showed transformation of black into orange scales. (D) The M3 eyespot was reduced in size, and the Cu1, Cu2, and A1 eyespots disappeared. On the dorsal forewing, the Cu1 eyespot showed transformation of black into orange scales. (E) The M1 eyespot on the right forewing was reduced in size, and the Cu1 eyespot showed transformation of black into orange scales. The chevron pattern on the wing margin and the central symmetry system bands running the length of each wing were distorted. (F) The M2, M3, and Cu1 eyespots on the left hindwing showed transformation of black into orange scales. (G) The M1 eyespot on the left forewing disappeared, and the Cu1 eyespot showed transformation of orange into black scales. Mutated eyespots are marked with red arrowheads. Scale; 5 mm.

**Fig. S7.**
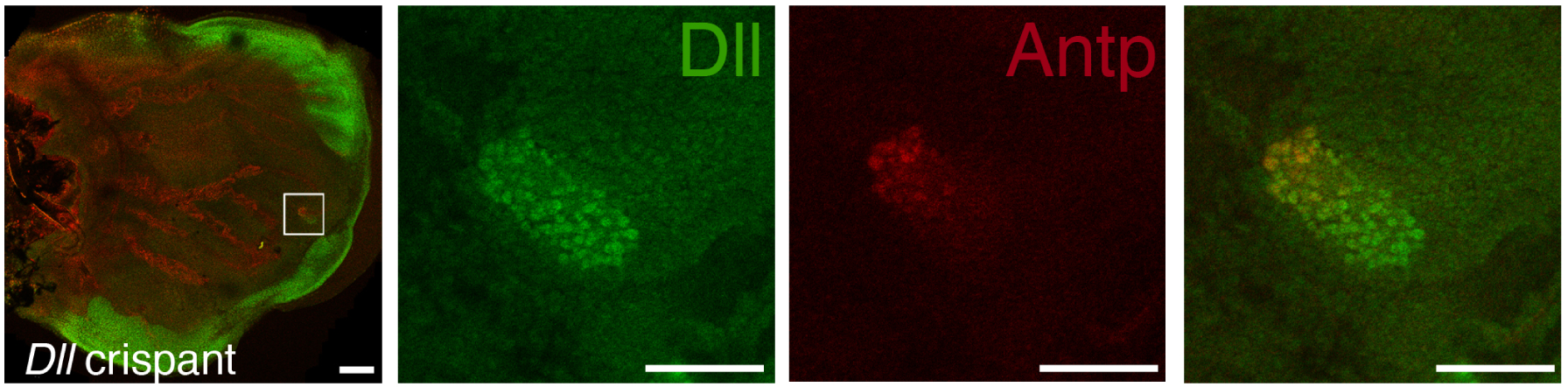
Expression pattern of Dll and Antp proteins in a *Dll* crispant Antp expression was only observed within the *Dll*-positive cells. Scale; 100 μm for low magnification and 50 μm for high magnification.

**Fig. S8.**
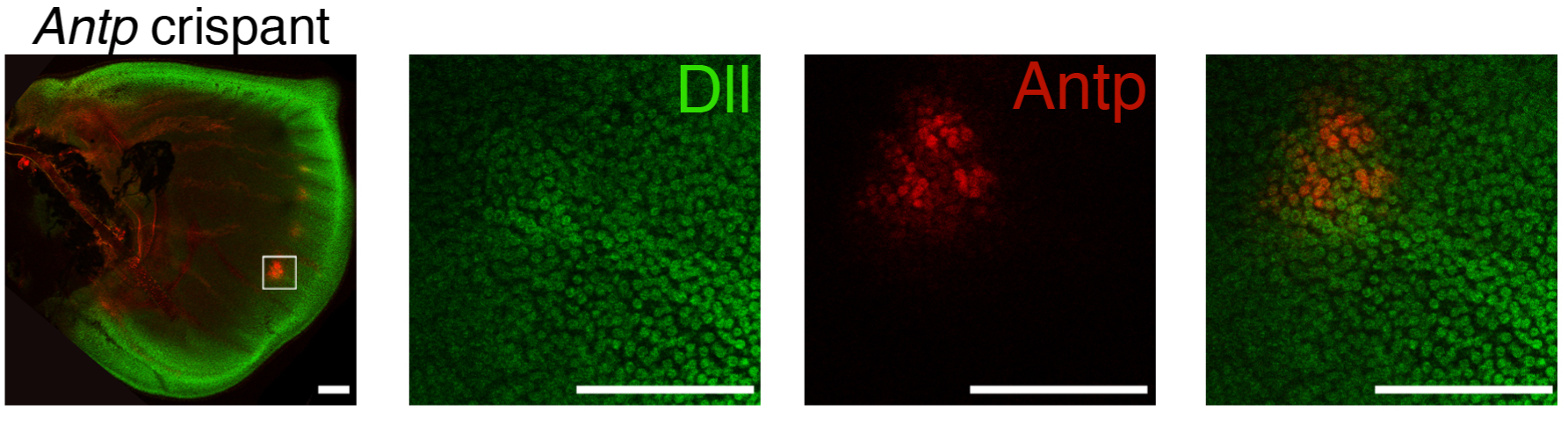
Expression pattern of Dll and Antp proteins in an *Antp* crispant. (A) Antp expression in Cu1 eyespot was partially lost, but Dll expression was not affected. Scale; 100 μm for low magnification and 50 μm for high magnification.

**Fig. S9.**
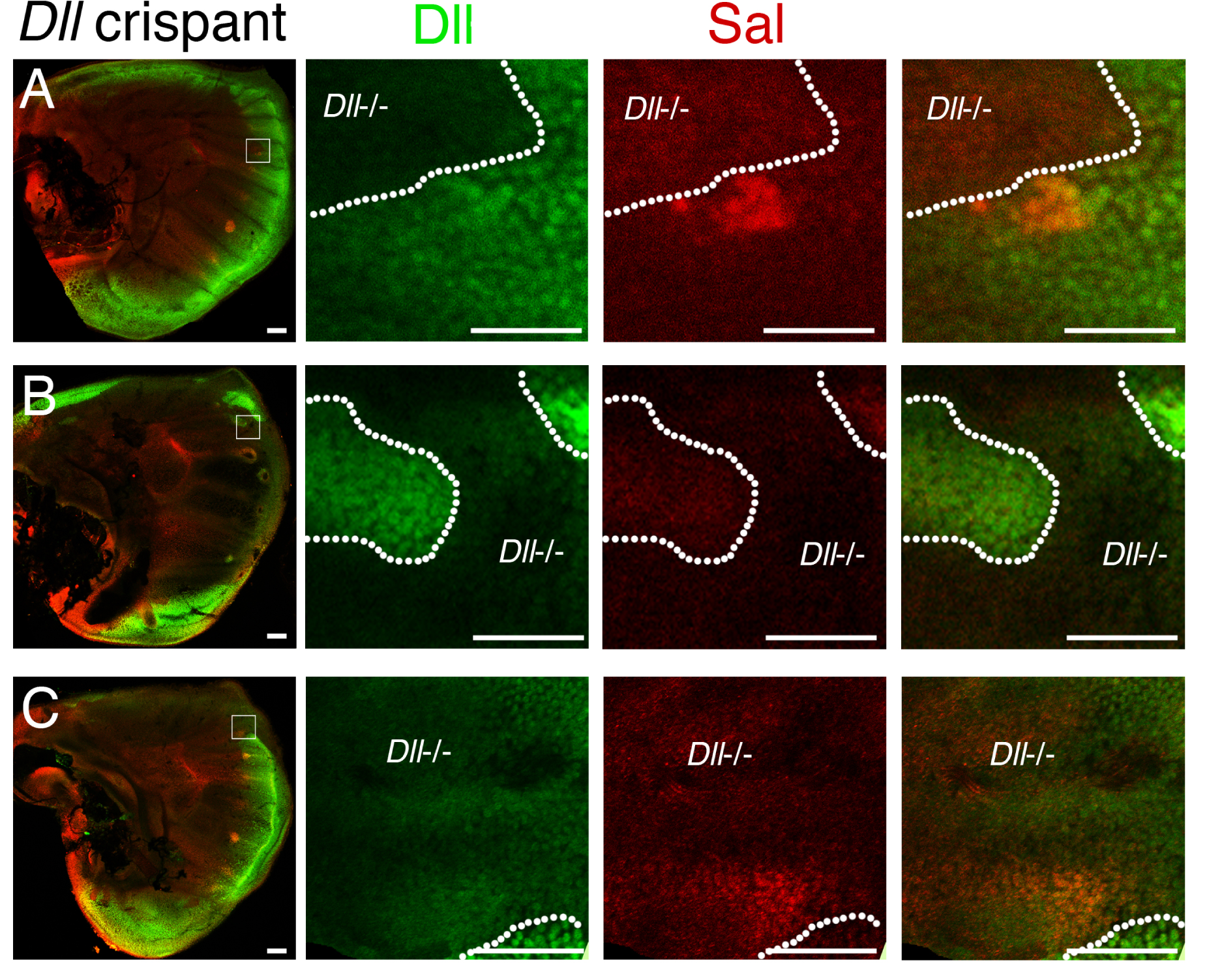
Expression pattern of Dll and Sal proteins in *Dll* crispants. (A) Sal expression was lost in *Dll* null mutant cells of the M1 eyespot. (B) Cells from the middle of the wing were broadly mutated and lost Sal activity. (C) Cells anterior to the M1 eyespot were mutated, which resulted in the loss of Sal expression in the eyespot centres. (D) Cells from the middle of the wing were broadly mutated, and in some wing sectors, Sal was ectopically expressed in the distal wing region. Scale; 100 μm for low magnification and 50 μm for high magnification.

**Fig. S10.**
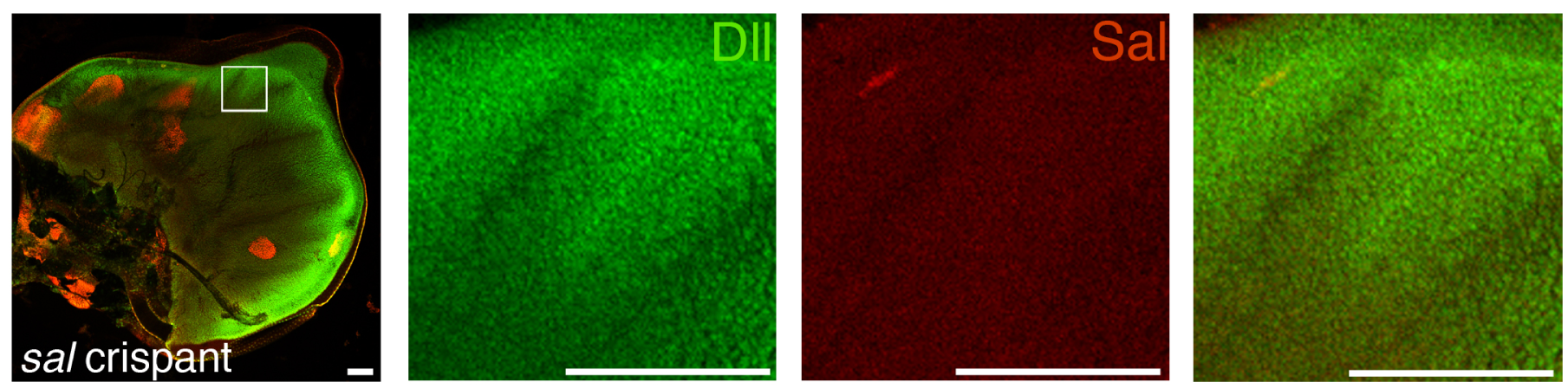
Expression pattern of Dll and Sal proteins in a *sal* crispant The distal wing region was broadly mutated for Sal activity, but Dll expression was not affected. Scale 50 μm.

**Fig. S11.**
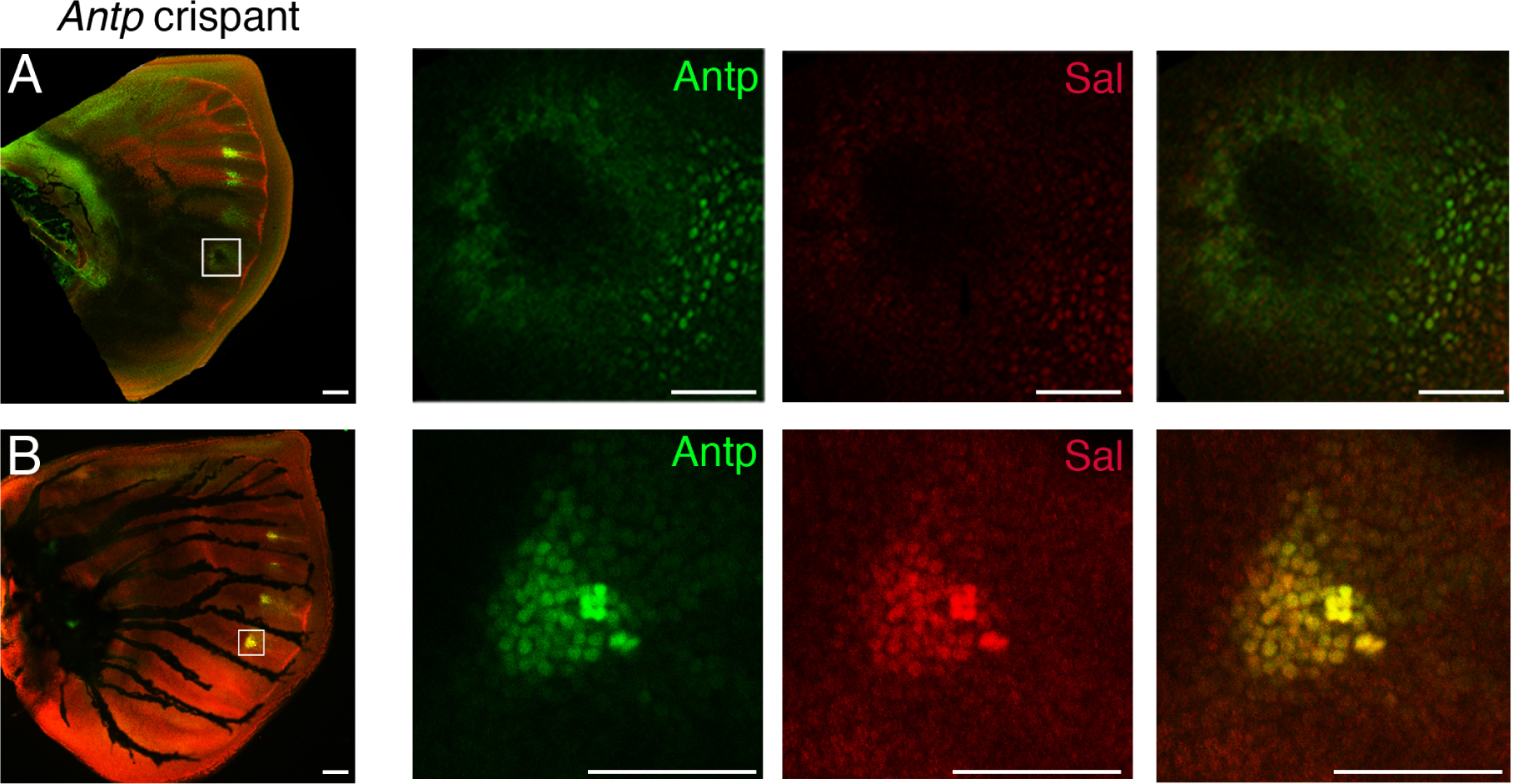
Expression pattern of Antp and Sal proteins in *Antp* crispants. (A) Sal expression was lost in *Antp* null mutant cells in the Cu1 eyespot. (B) Some Cu1 eyespot cells lost Antp activity, and Sal expression was only detected in the Antp-positive cells. Scale; 100 μm for low magnification and 50 μm for high magnification.

**Fig. S12.**
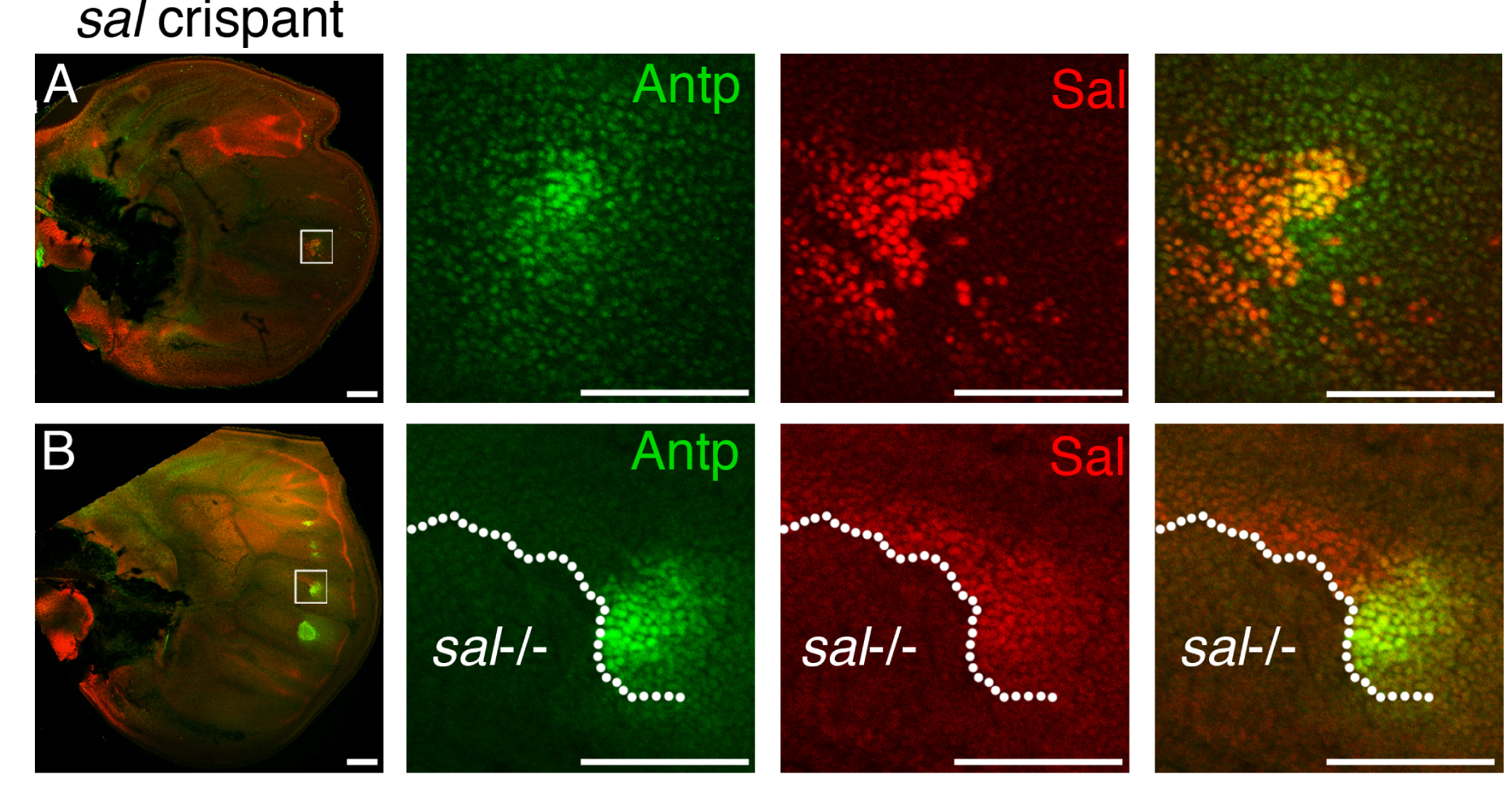
Expression pattern of Antp and Sal proteins in *sal* crispants. (A) The wing is broadly mutated for Sal activity. Some Cu1 eyespot cells lost Sal expression, and Antp expression was only detected in the Sal-positive cells. (B) Cells around the M3 eyespot were mutated, and Antp expression was detected only in the Sal-positive cells. Scale; 100 μm for low magnification and 50 μm for high magnification.

**Fig. S13.**
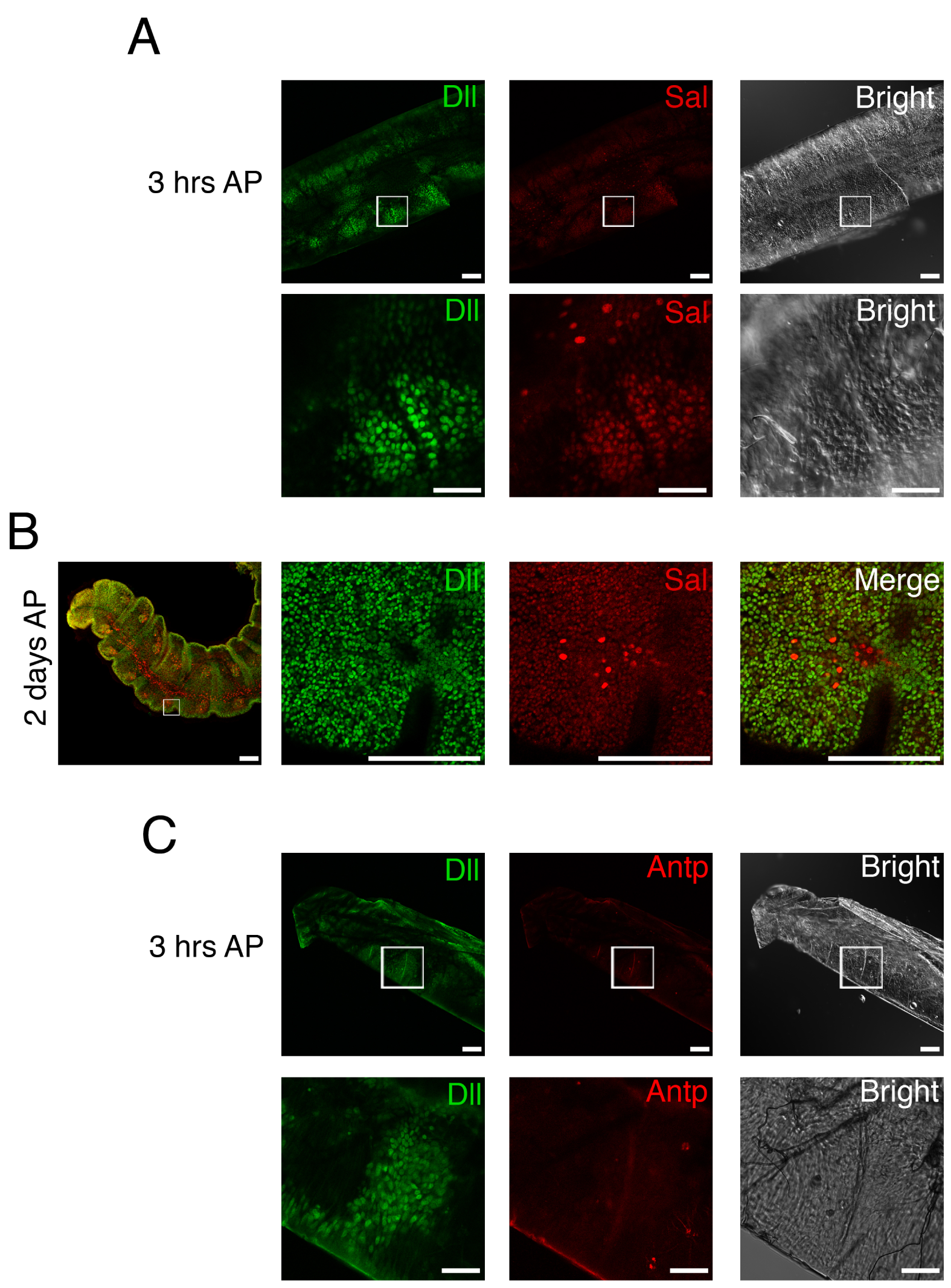
Expression of Dll, Sal, and Antp proteins in pupal antennae. (A) Dll and Sal were co-expressed in the segments of the developing antenna of a 3-h-old pupa. (B) Expression patterns of Dll and Sal in the pupal antenna, 2 days after pupation (AP). Dll was ubiquitously expressed in the antenna, but Sal expression is observed in the neurons. (C) Expression patterns of Dll and Antp in the antenna of a 3-h-hold pupa. Antp expression was not observed in the antenna. The regions within the white squares are shown at higher magnification. Scale; 100 μm for low magnification and 50 μm for high magnification.

**Fig. S14.**
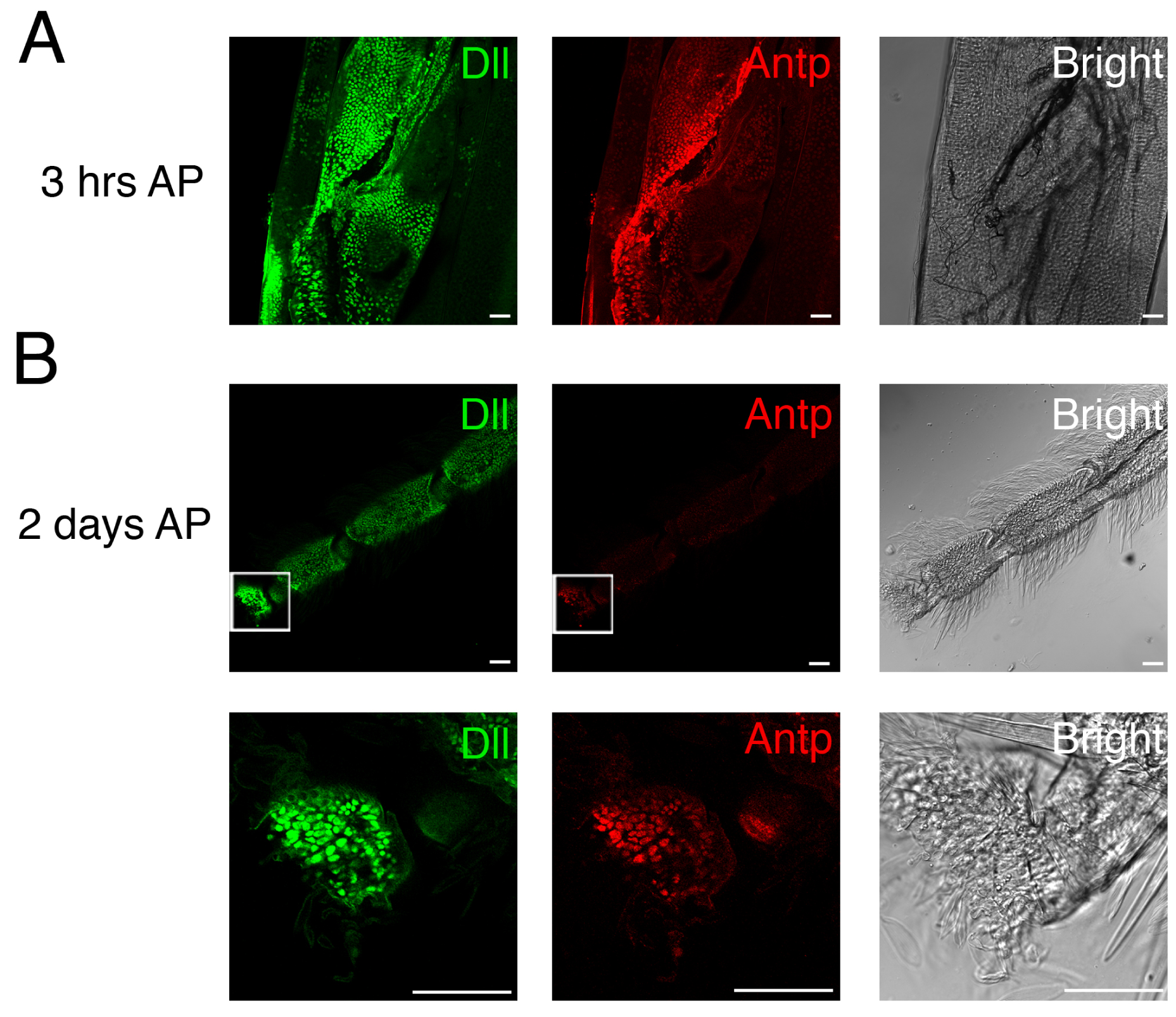
Expression patterns of Dll, and Antp proteins in pupal legs. (A) Expression pattern of Dll and Antp in the developing leg of a 3 -h-old pupa. Dll and Antp are co-localized in the developing pupal leg. (B) Expression patterns of Dll and Antp in the leg of a 3-days-old pupa. Dll is ubiquitously expressed in the distal region of a developing pupal leg. Antp is also expressed in the developing pupal leg. The regions within the white squares are shown at higher magnification. Scale; 100 μm.

**Fig. S15.**
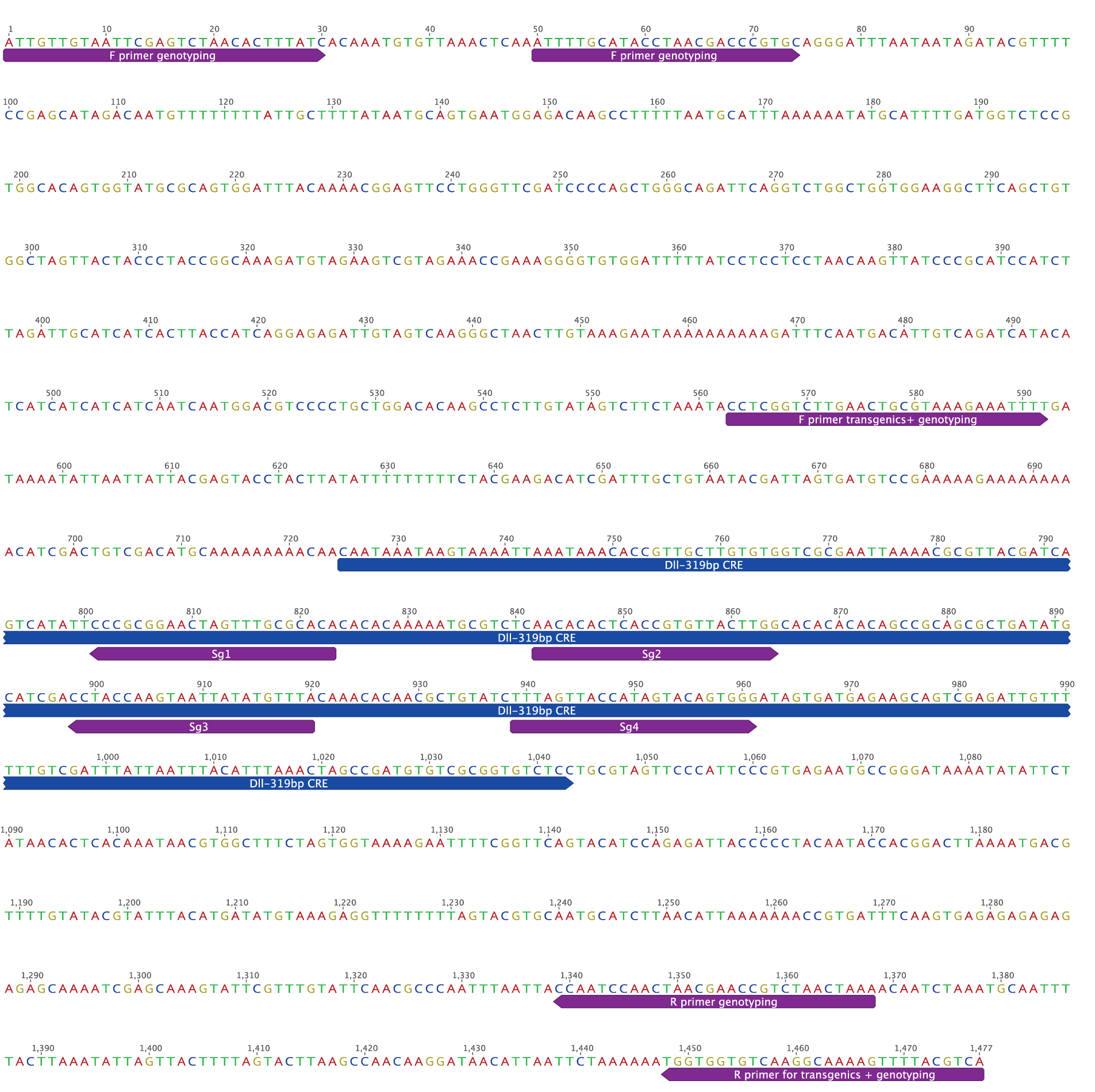
*Dll319* CRE annotated with CRISPR guides and primers used for the transgenics and genotyping crispants.

**Fig. S16.**
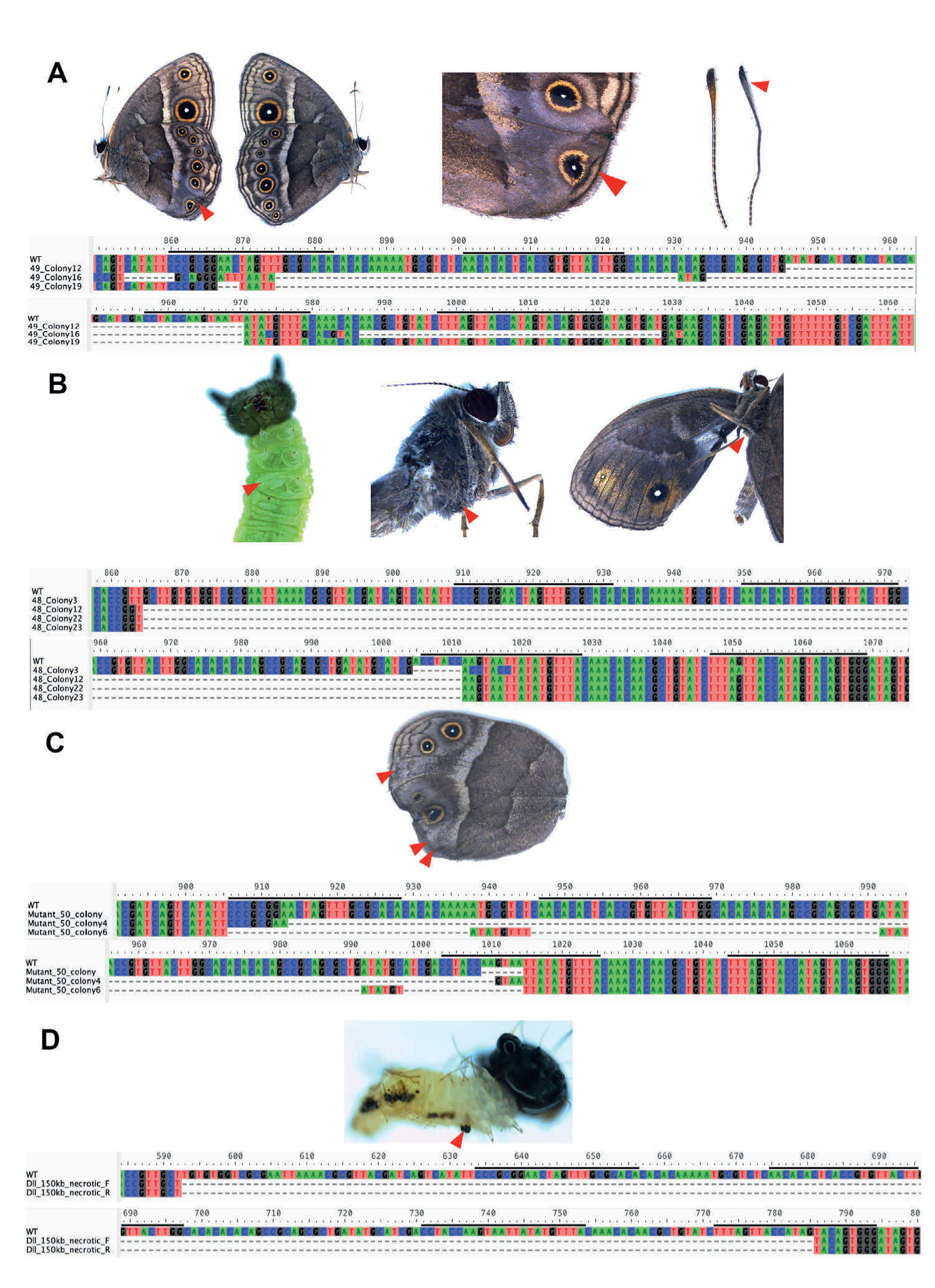
Sanger sequencing results following CRISPR-targeting of the *Dll319* CRE in four crispant individuals, WT refers to the wild-type sequence. Positions of the CRISPR guides are shown as horizonal black lines above each sequence. (A) Colony PCR sequencing results with variable sized deletions for a crispant with a missing eyespot; pigmentation defects are visible on the wing and antenna. The left image shows the wing defects. The mirror image on the right shows wild-type wing phenotype. (B) The same individual from Fig. 3C is shown with a missing T3 leg in the larva and adult, as well as a missing wing. The sequence of a colony PCR product revealed a 147 bp deletion. (C) Colony PCR sequencing result showing a 108 bp deletion for an individual with three missing eyespots. (D) Sanger sequencing showing a 193 bp deletion from a whole larva that showed areas of necrosis and a missing distal tip of a T3 leg.

**Fig. S17.**
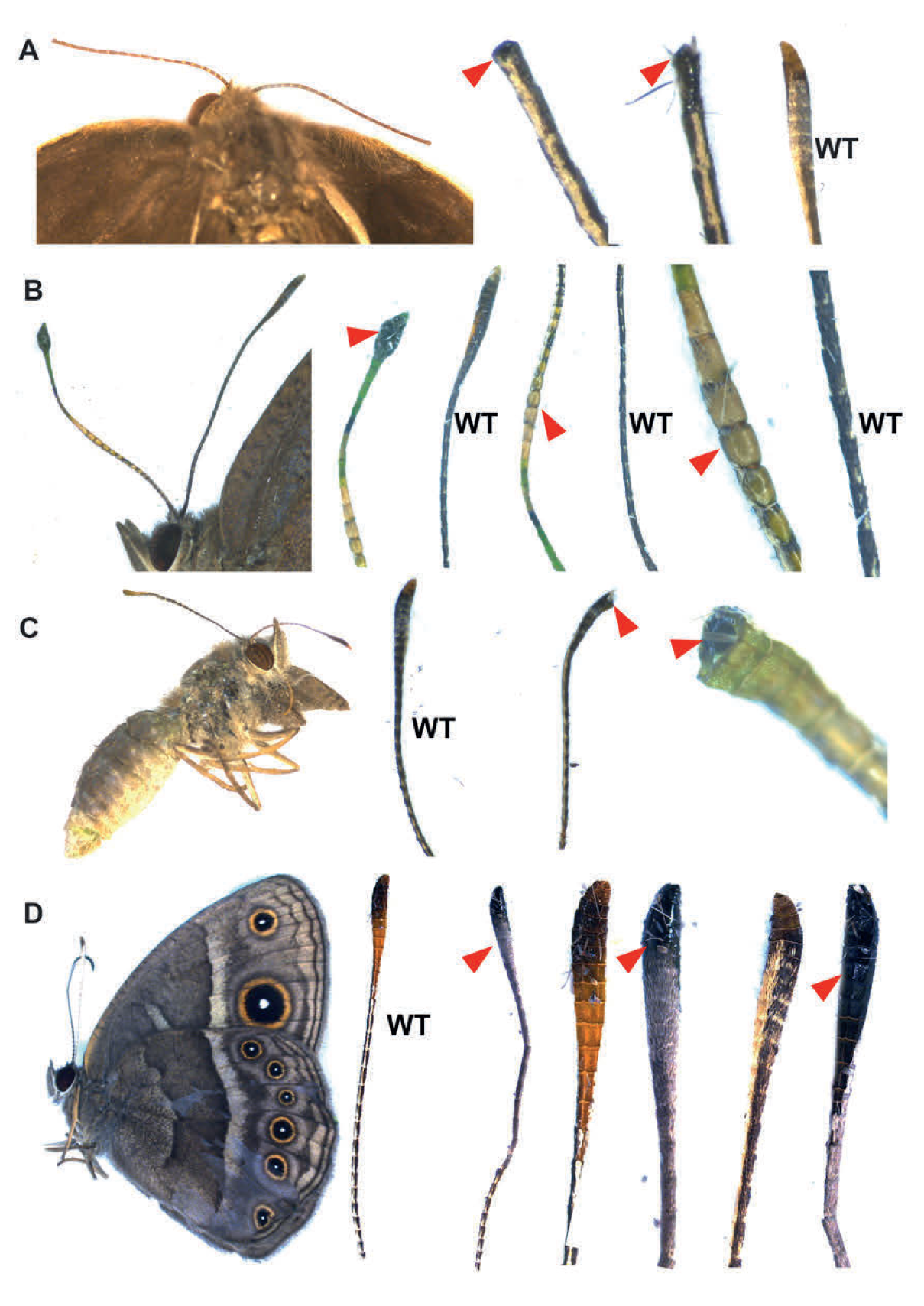
Four individual crispants showing antennal deformities following disruption of the *Dll319* CRE. (A) Both antennae of this butterfly were missing the distal tip. (B) One antenna showed a developmental abnormality, changing the shape of the distal tip. Additionally, the stem of the antenna had a different morphology to the wild-type and also showed loss of scales. (C) The very distal tip of one antenna in this individual was missing. (D) One antenna was crooked and showed a change in pigmentation from brown to grey. Furthermore, we noted a loss of scales on one side of the antenna that has been replaced by a shiny black cuticle, as compared to wild-type. Red arrows point to the antennae with developmental defects.

**Fig. S18.**
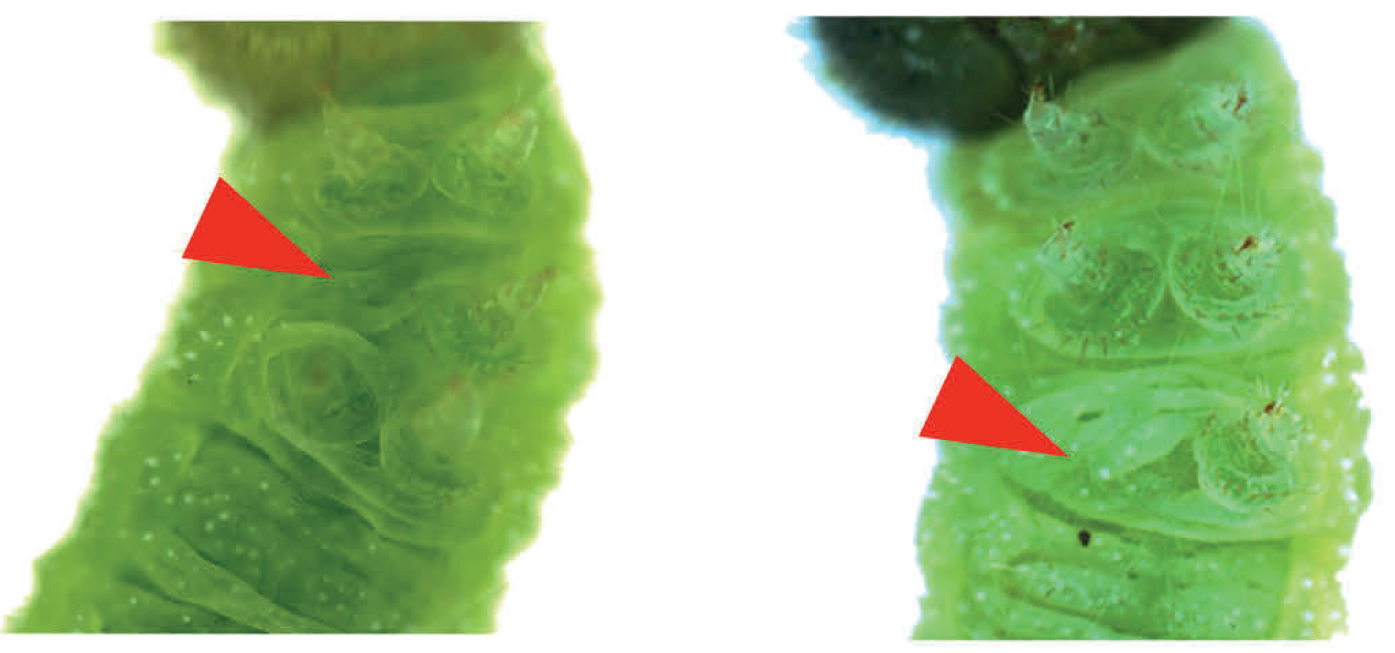
Targeting of the *Dll319* CRE led to losses of either T2 or T3 legs.

**Fig. S19.**
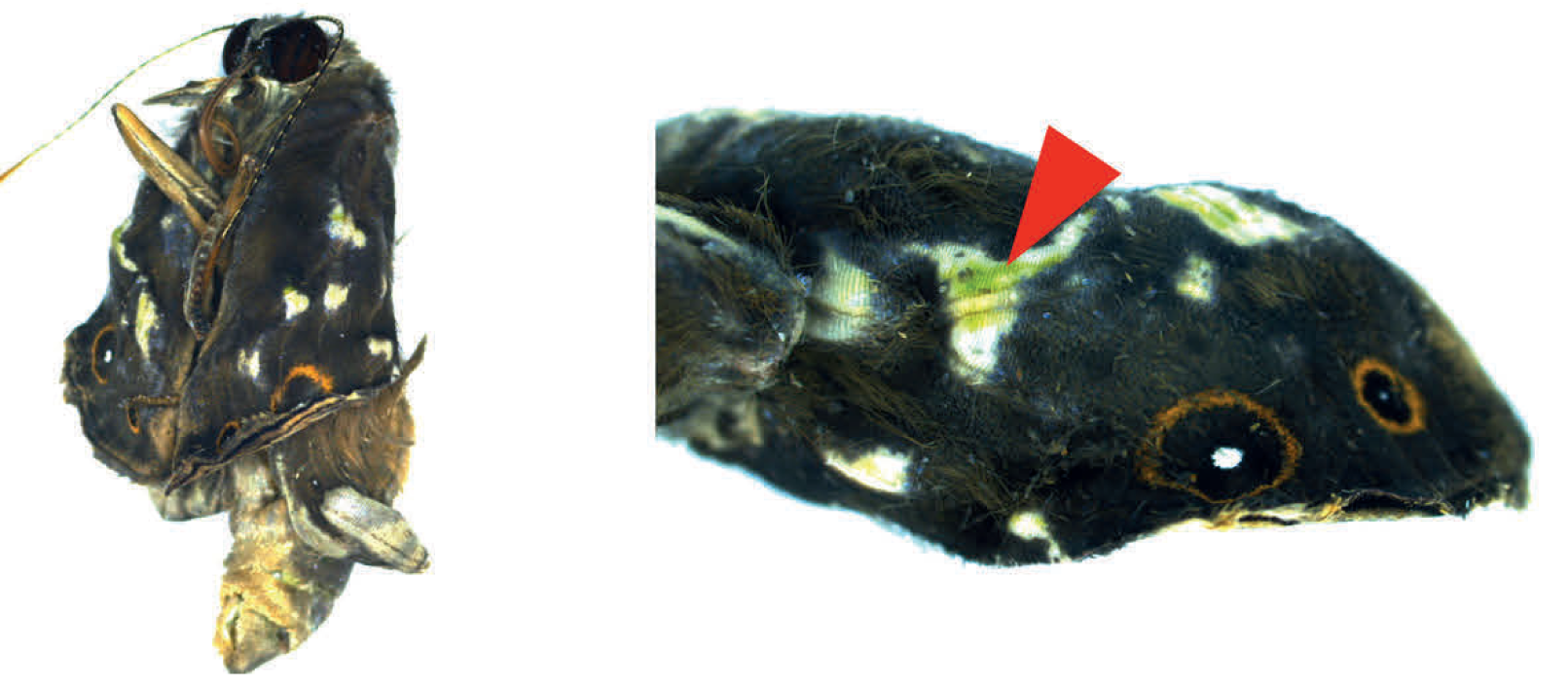
A *Dll319* crispant shows white wing patches due to a complete loss of wing scales.

**Fig. S20.**
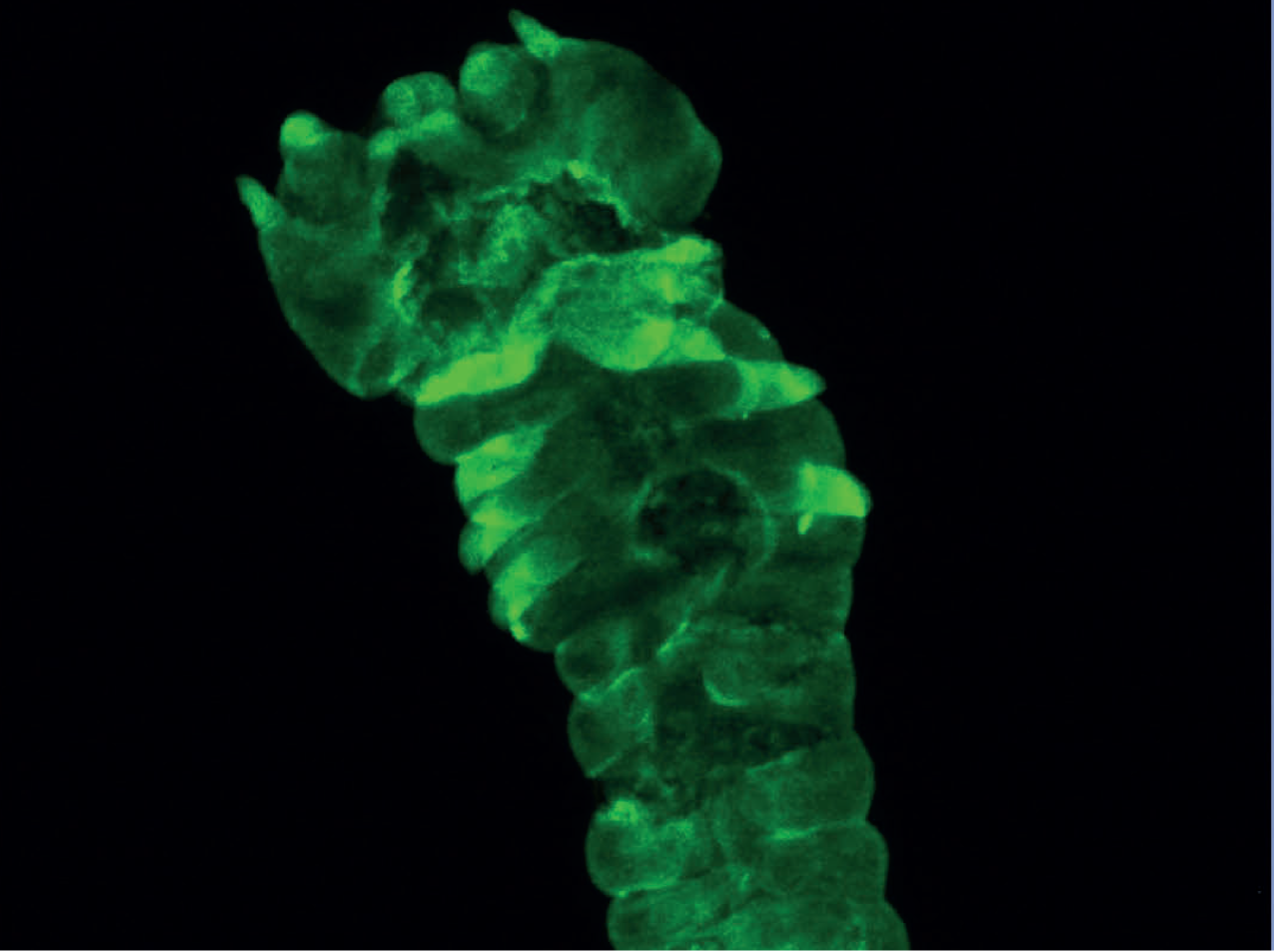
Second replicate of an EGFP-expressing embryo from the F3 transgenic generation, where the *EGFP* gene was driven by the *Dll319* CRE. EGFP was visualized through the use of anti-EGFP antibodies.

**Fig. S21:**
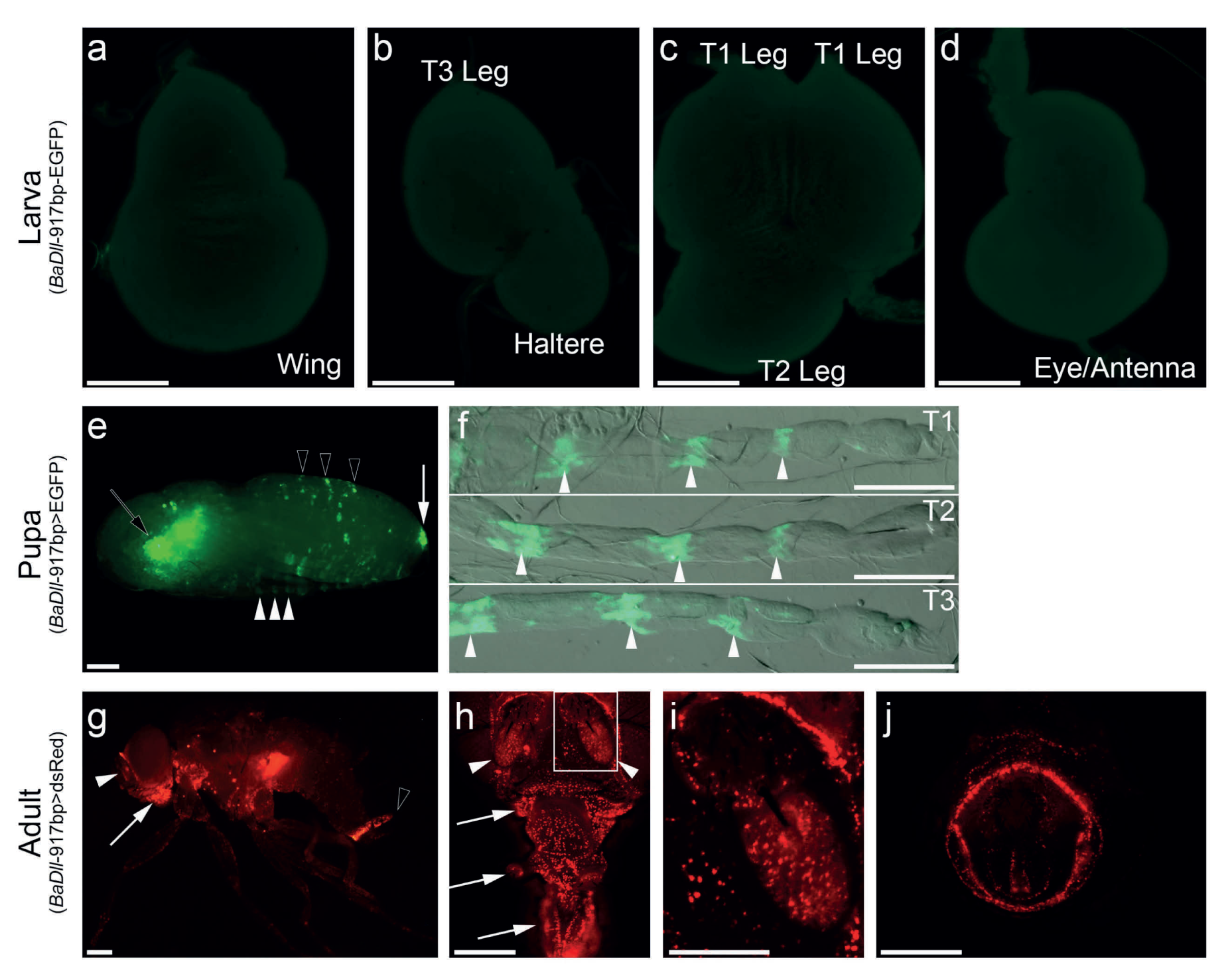
Cross-species reporter assay of *Dll319* CRE in *D. melanogaster*. **a-d**. Imaginal discs of *Dll319*-EGFP flies. No noticeable enhancer activity is detected at this stage. **e,f**. *Dll319* CRE activity during the pupal stage as visualized by *Dll319*-Gal4>UAS-EGFP. **e**. Enhancer activity is detected in the pupal legs (white arrowheads), genitalia (white arrow), and in the abdomen (black arrowheads). EGFP in the eye (black arrow) is transgenic marker expression. **f**. Close-up images of the pupal legs, showing the enhancer activity in the tarsal segments (white arrowheads). **g-j**. *Dll319* CRE activity at the adult stage visualized by *Dll319*-Gal4>UAS-dsRed. **g**. *Dll319* CRE drives low ubiquitous ectodermal expression, with increased expression in the adult antennae (white arrowhead), mouthparts (white arrow), and genitalia (black arrowhead). **h**. Close-up of adult head, showing the increased activity in antennae (white arrowheads) and mouthparts (white arrows). **i**. Close-up of antenna enhancer activity within the area indicated by the white box in **h**. **j**. Close-up of enhancer activity in the adult genitalia. A lack of enhancer activity during the last larval stage may be due to the nature of this enhancer or a limitation of testing the activity of this enhancer in a cross-species setting (or a combination of both). Nonetheless, the presence of increased enhancer activity in multiple tissues, especially with these that are homologous to antennae, suggests that *Dll319* contains a pleiotropic CRE. Scale bars indicate 100 µm (**a-d**, **f**, **i**) and 200 µm (**e**, **g**, **h**, **j**).

**Fig. S22.**
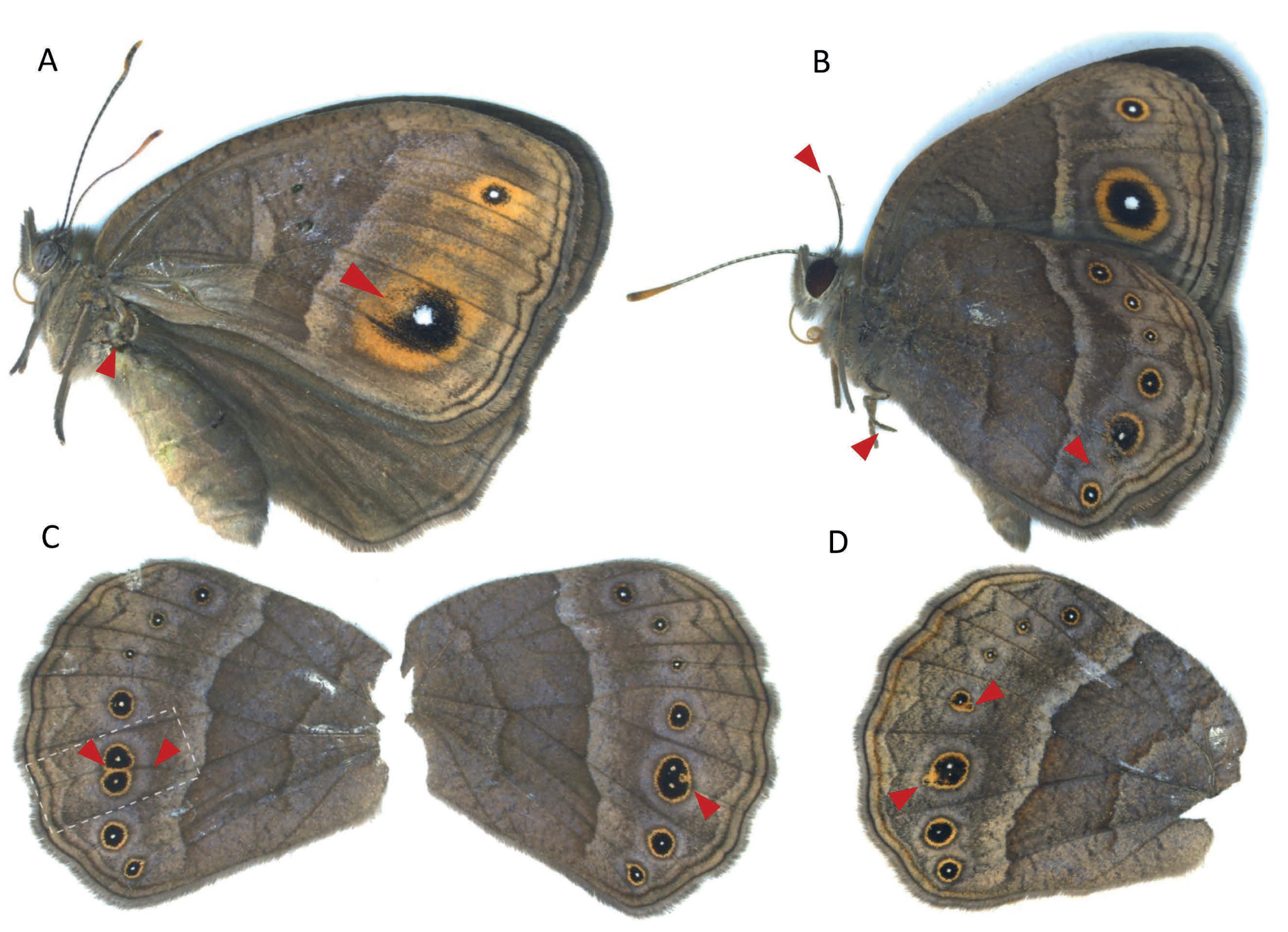
*sal* CRE (*sal*740) crispants phenotypes. A. Loss of hindwing and black pigment in the forewing. B. Loss of eyespot in Cu2 hindwing sector, loss of distal part of the antenna and crooked T3-leg. C. Left wing: Additional vein formation in Cu1 sector of hindwing with split of Cu1 eyespot. Right wing: No ectopic vein but also split of Cu1 eyespot showing two white centres and another eyespot pigment deformity. D. Ectopic eyespot in Cu1 and M3 sector in hindwing.

**Fig. S23.**
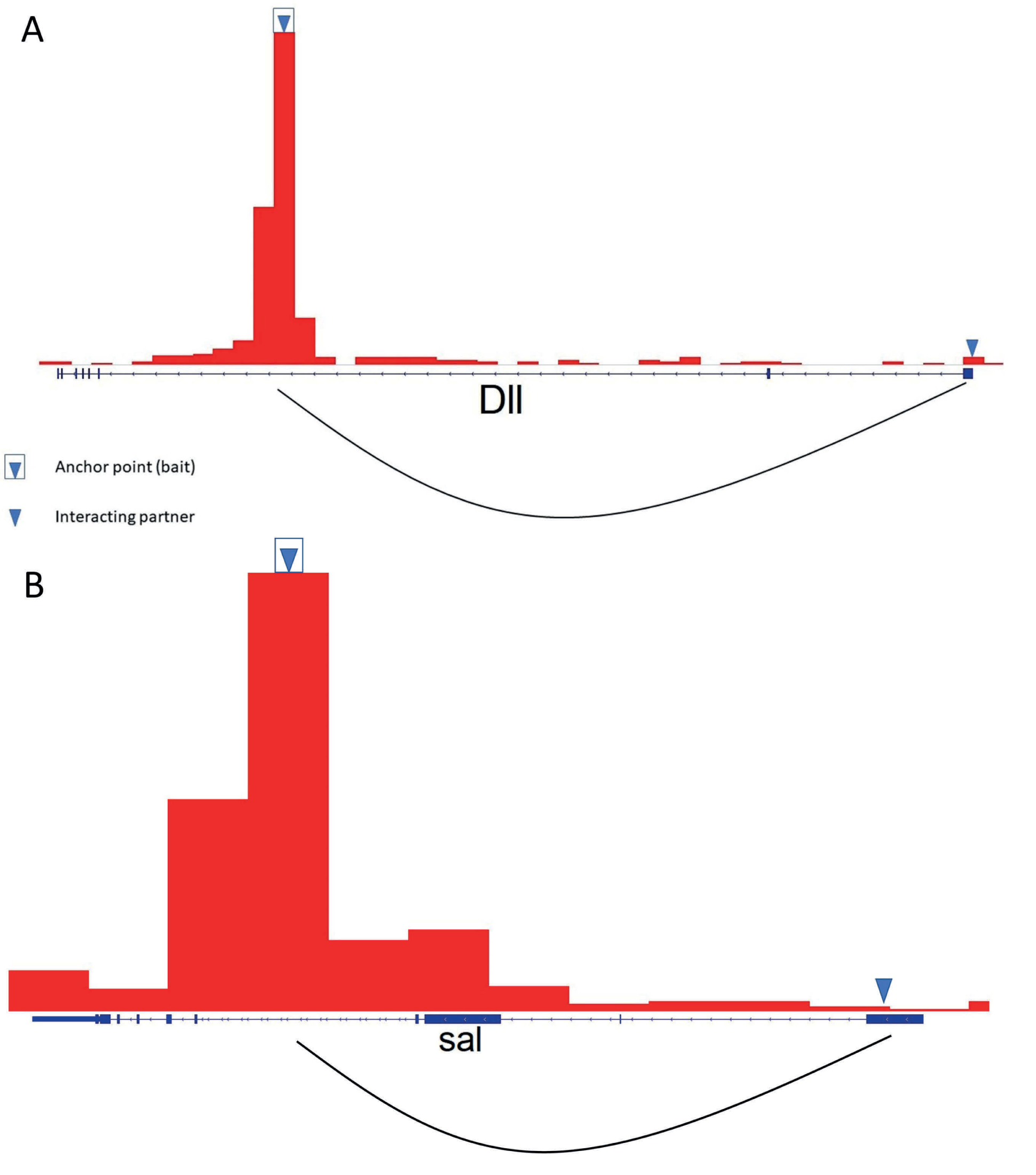
Virtual 4c plot for *Dll319* and *sal740.* (A) The graph around *Dll319* region showing its interaction with Dll promoter region. (B) Graph around *sal740* region shows its interaction with sal promoter region

**Fig. S24.**
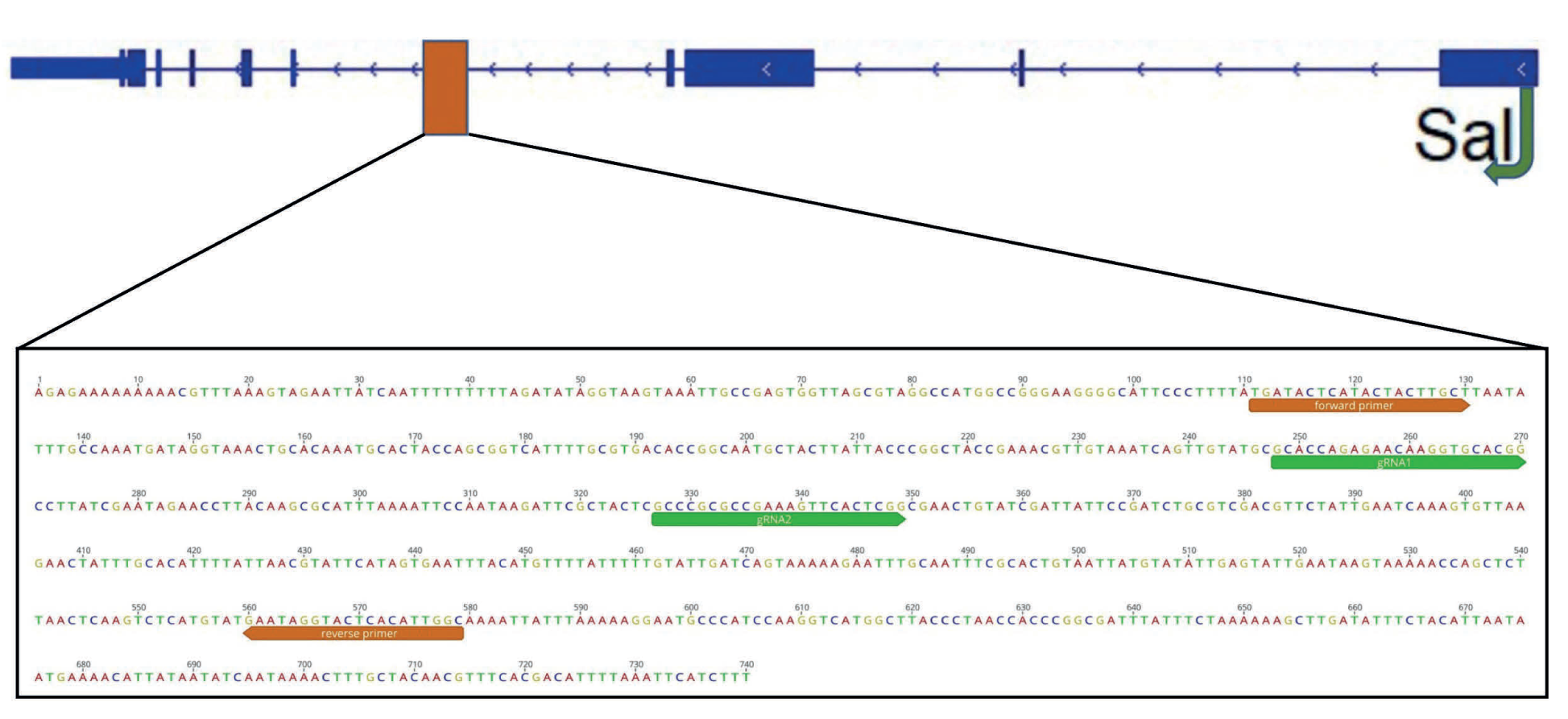
*sal740* CRE region annotated with CRISPR guides and primers used for knockout and genotyping

**Fig. S25.**
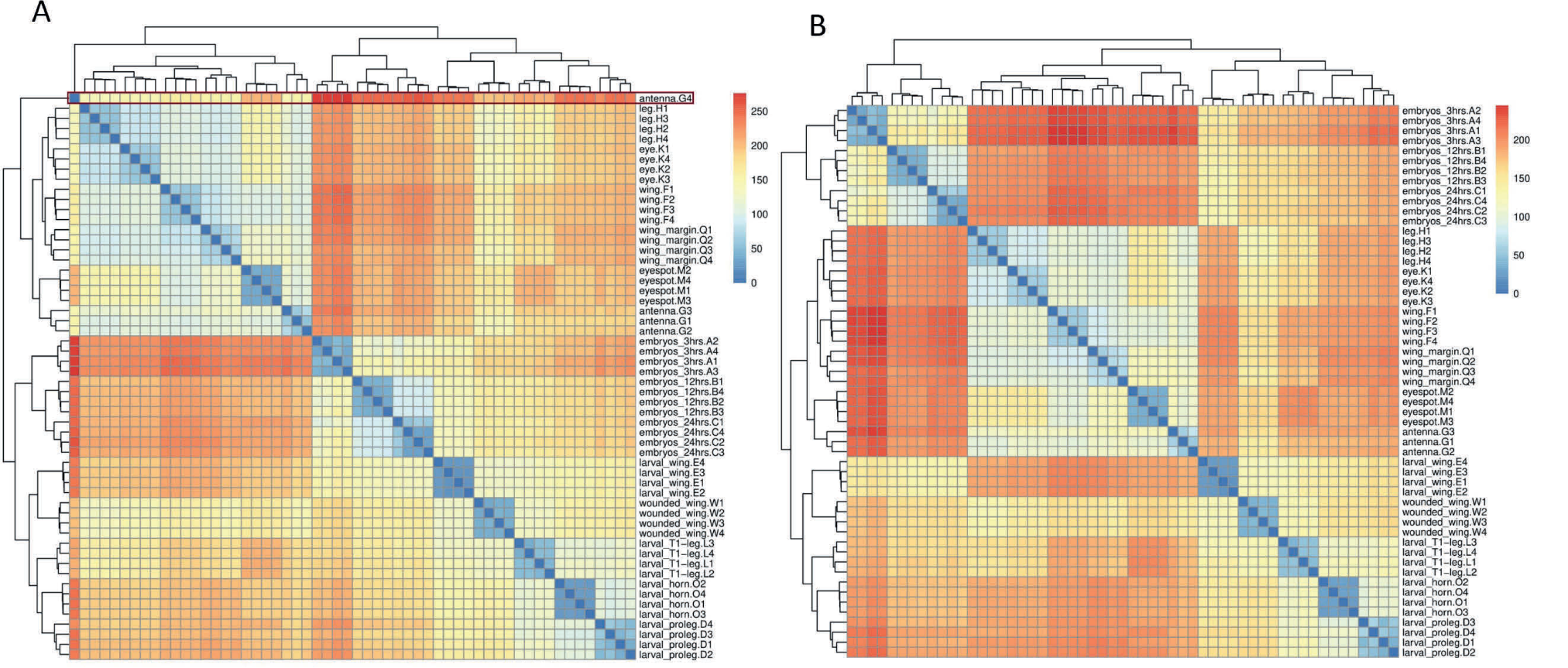
Sample clustering using an Euclidean distance matrix before and after sample filtering. (A) Initial sample clustering shows antenna. G4 is an outlier. (B) After excluding the outlier (antenna.G4), the replicates from each group cluster together.

**Fig. S26.**
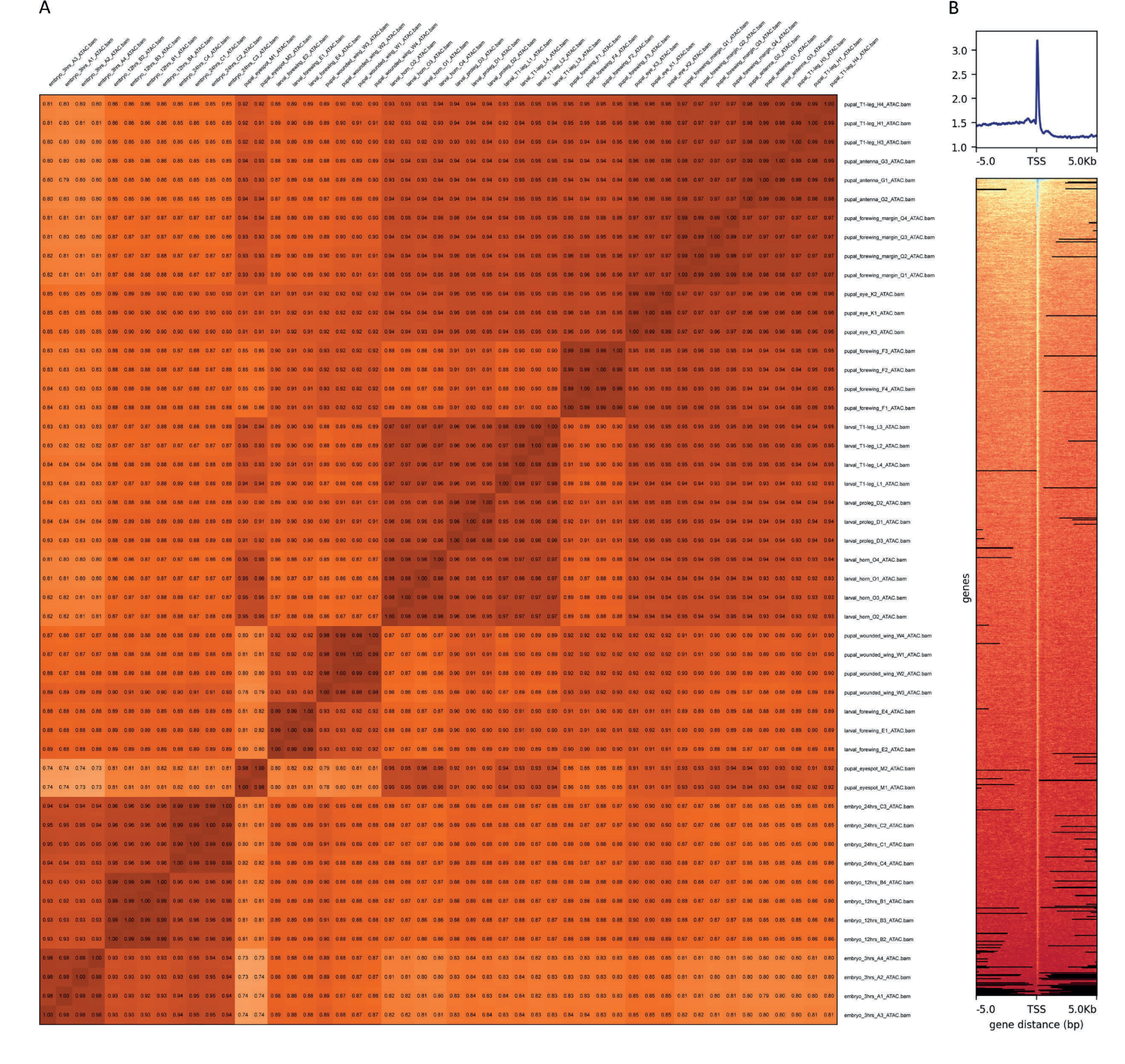
Quality assessment of ATAC libraries. A. Correlation matrix between the replicates in the group. B. ATAC peaks are highly enriched in Transcription start sites (TSS) and this highlights libraries of good quality.

## Supplementary Tables

**Table S1.** Number of injected individuals that displayed developmental defects due to CRISPR-targeting of the *Dll319* CRE.

| No. eggs injected | No. hatched | Survival 3<sup>rd</sup> instar | Leg mutants | Missing eyespots | Deformed antennae | Missing wings | Pigment defects |
| --- | --- | --- | --- | --- | --- | --- | --- |
| 366 | 40 | 35 | 1 | 0 | 0 | 0 | 0 |
| 646 | 56 | 51 | 1 | 2 | 0 | 1 | 3 |
| 422 | 87 | 76 | 1 | 2 | 1 | 0 | 4 |
| 797 | 193 | 186 | 1 | 0 | 4 | 0 | 0 |

**Table S2.**
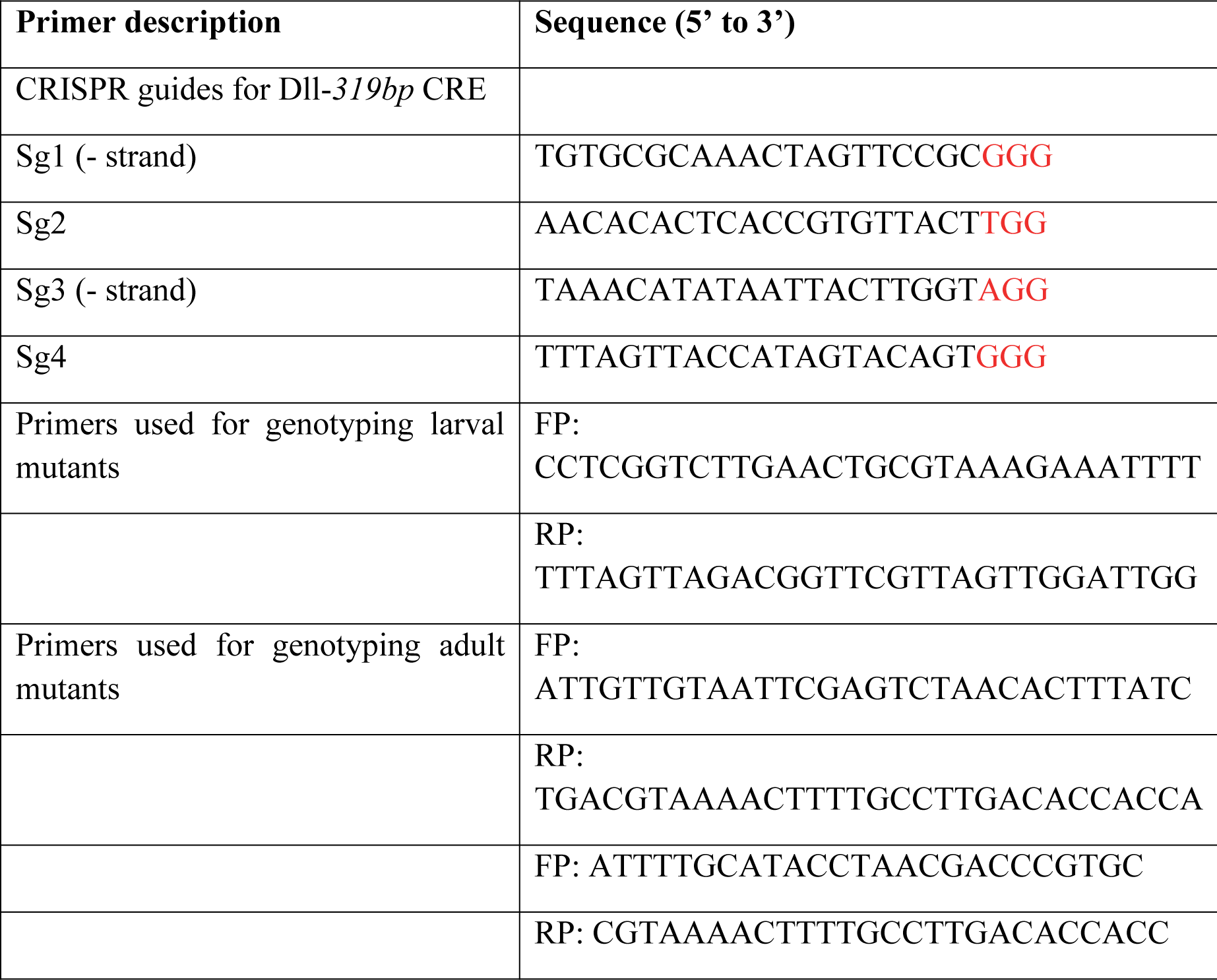
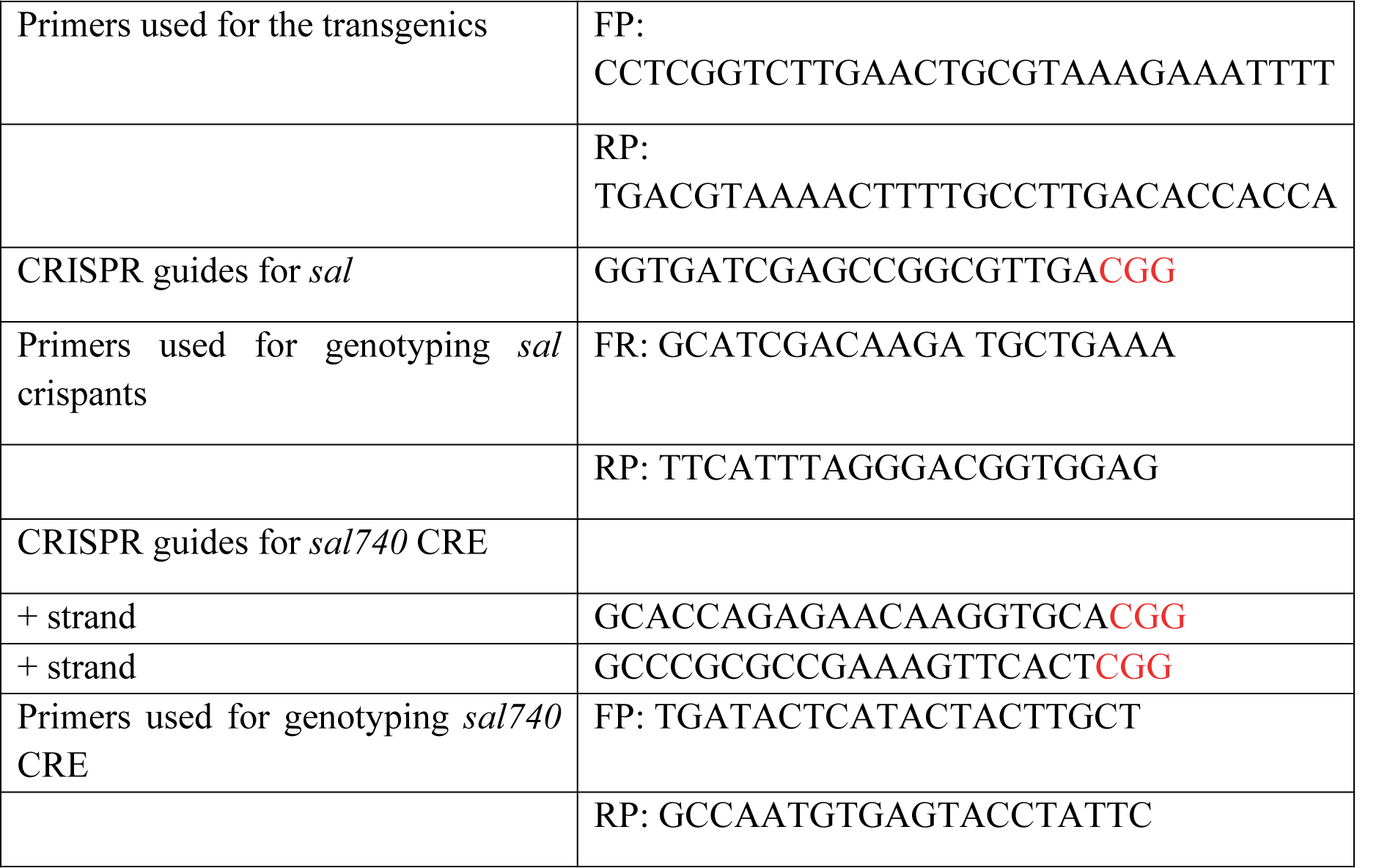
Primers used for CRISPR guide RNAs and for genotyping crispants.

**Table S3.** Number of injected individuals that displayed developmental defects due to CRISPR-targeting of *sal*.

| No. eggs injected | No. hatched | Survival 3<sup>rd</sup> instar | No. surviving until adulthood | Number of adults showing phenotype |
| --- | --- | --- | --- | --- |
| 108 | N.A. | N.A. | 51 | 11 |
| 102 | N.A. | N.A. | 23 | 5 |

**Table S4.** Number of injected individuals that displayed developmental defects due to CRISPR-targeting of the *sal740* CRE

| No. eggs injected | No. hatched | Survival 3<sup>rd</sup> instar | Leg mutants | Eyespot mutant | Deformed antennae | Wing mutant | Horn defect | Vein defect |
| --- | --- | --- | --- | --- | --- | --- | --- | --- |
| 174 | 23 | 17 | 0 | 2 | 1 | 0 | 1 | 1 |
| 116 | 27 | 21 | 1 | 4 | 2 | 1 | 0 | 1 |

**Table S5.** Tissue types and number of individuals used for each replicate in RNA-seq

| Tissue | Number of individuals |
| --- | --- |
| Cu1 control tissue – Nes1 | 20 |
| Cu1 eyespot | 20 |
| M3 control tissue - Nes2 | 20 |
| Wings (larval, pupal 3 h, wounded 24 h) | 15 |
| Embryos (3 h, 12 h, 24 h) | 15 |
| T1- legs (larval, pupal 3 h) | 15 |
| Pupal antennae (3 h) | 15 |
| Pupal eyes (3 h) | 15 |
| Pupal wing margins | 15 |
| Larval prolegs | 15 |
| Larval horns | 15 |

Dissections were performed from the right and left sides of each animal, except for wounded wings, where only one side of each pupa was wounded.

**Table S6.**
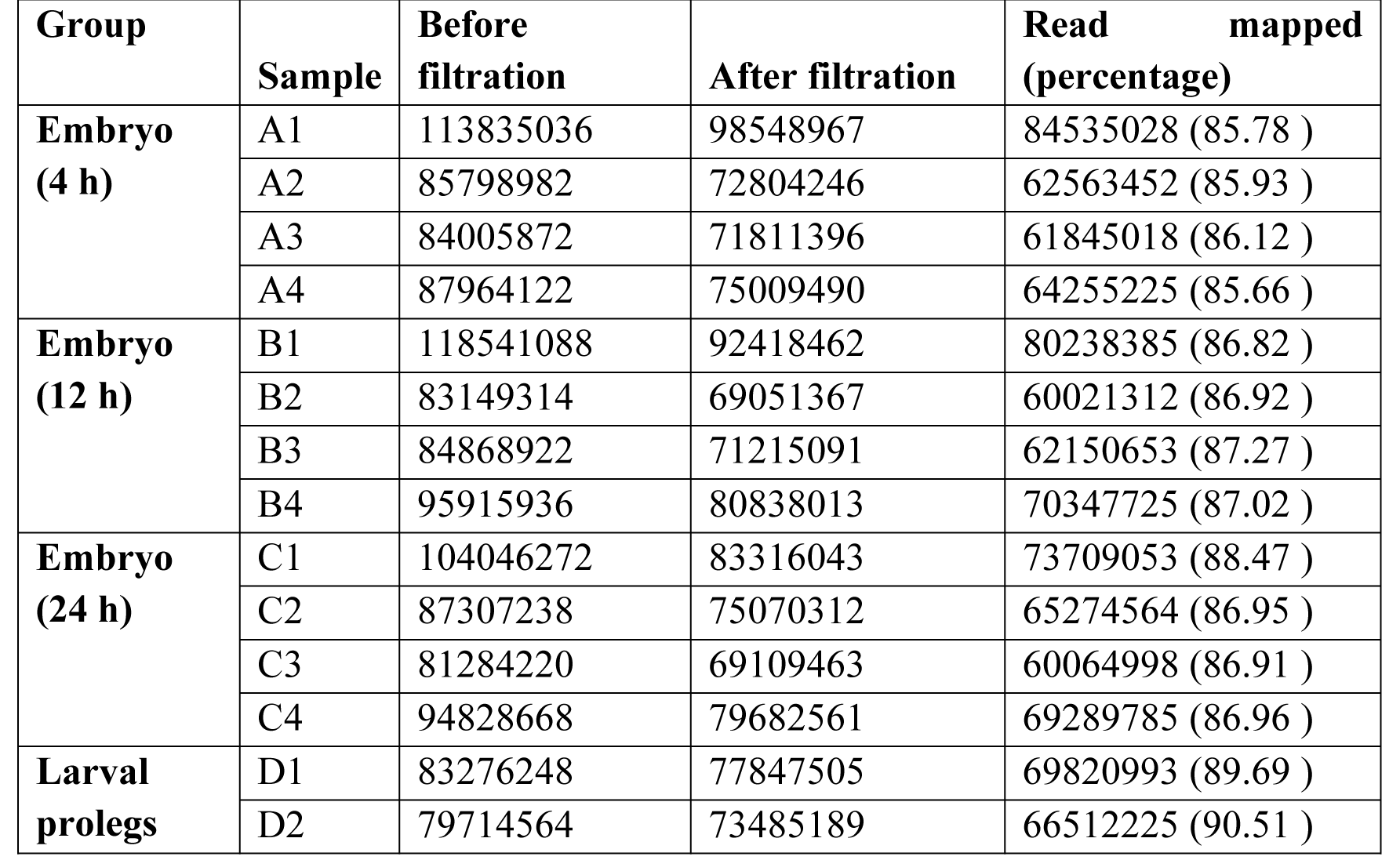
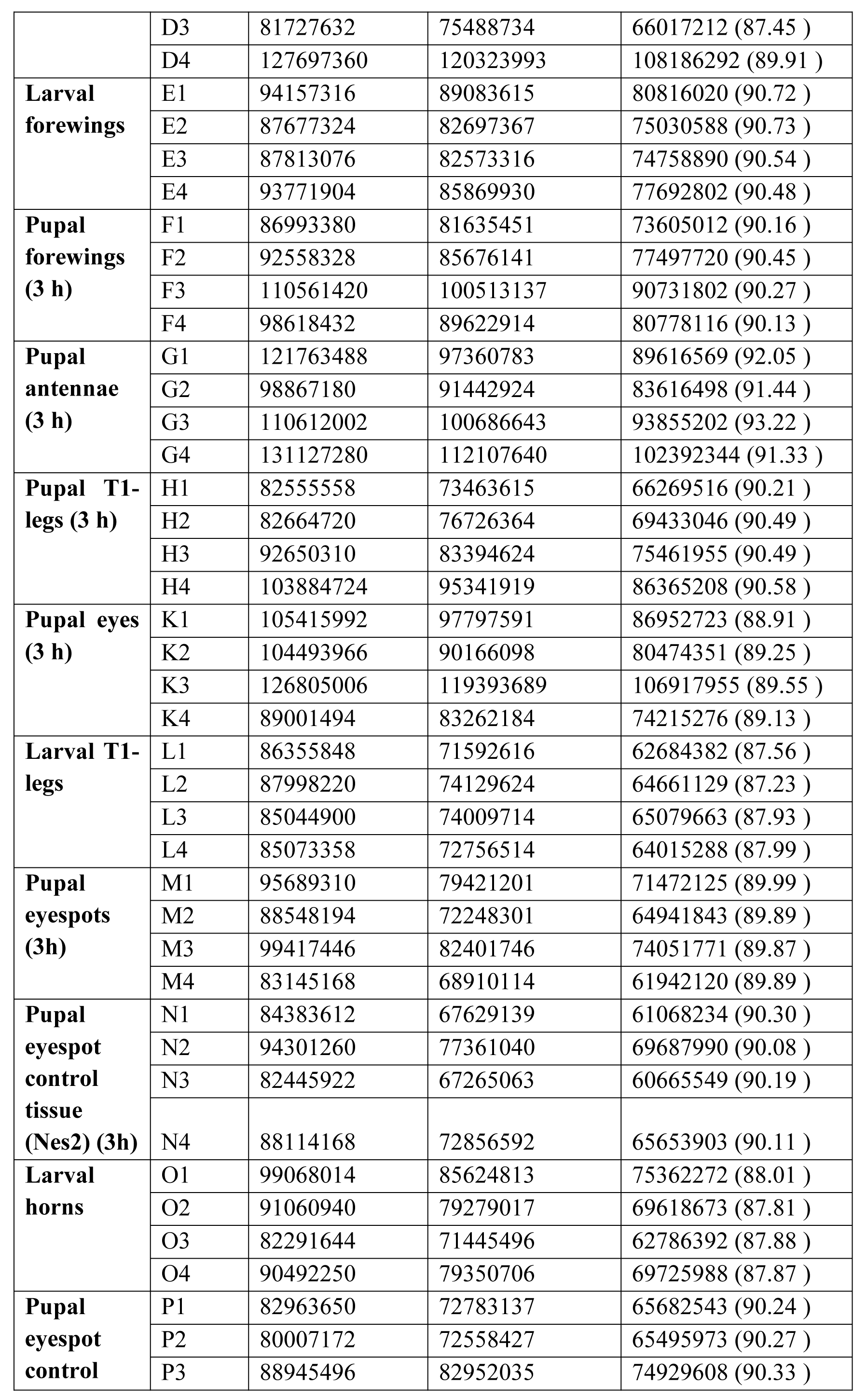
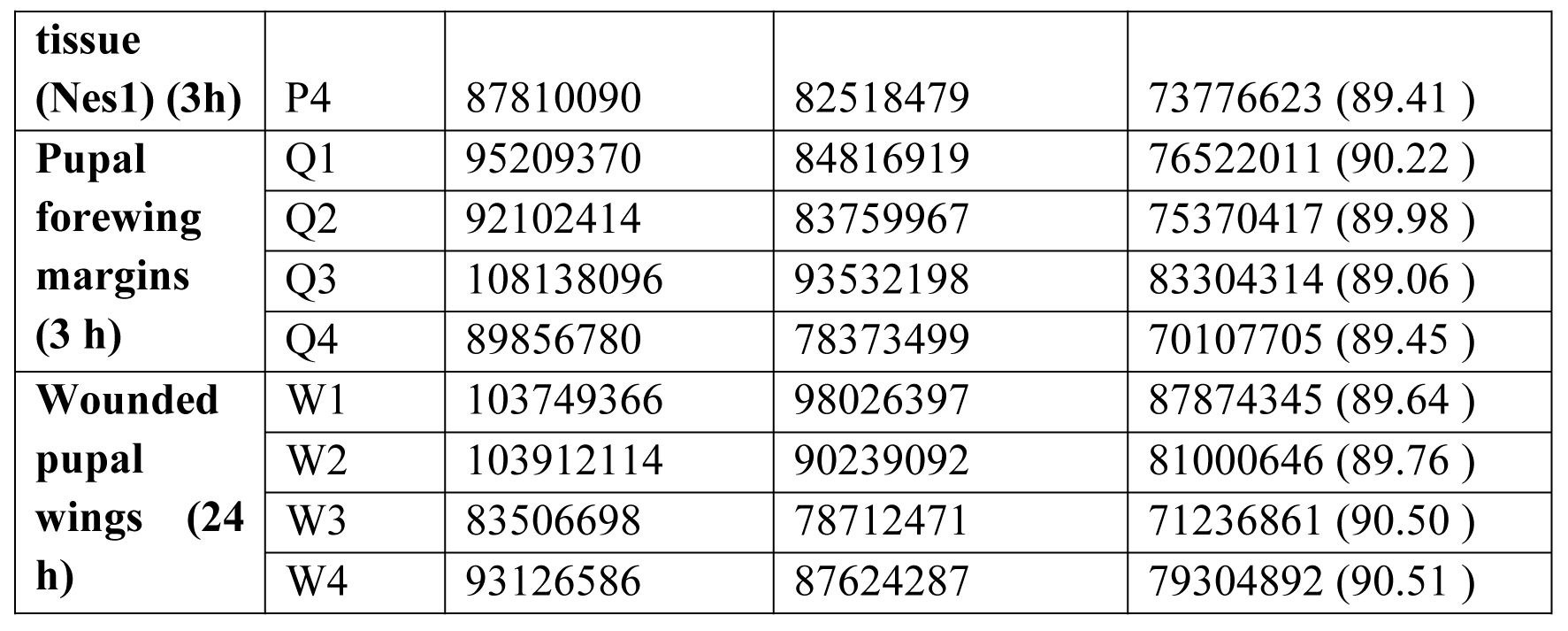
RNA sequencing data. Read-depth and alignment rate

**Table S7.**
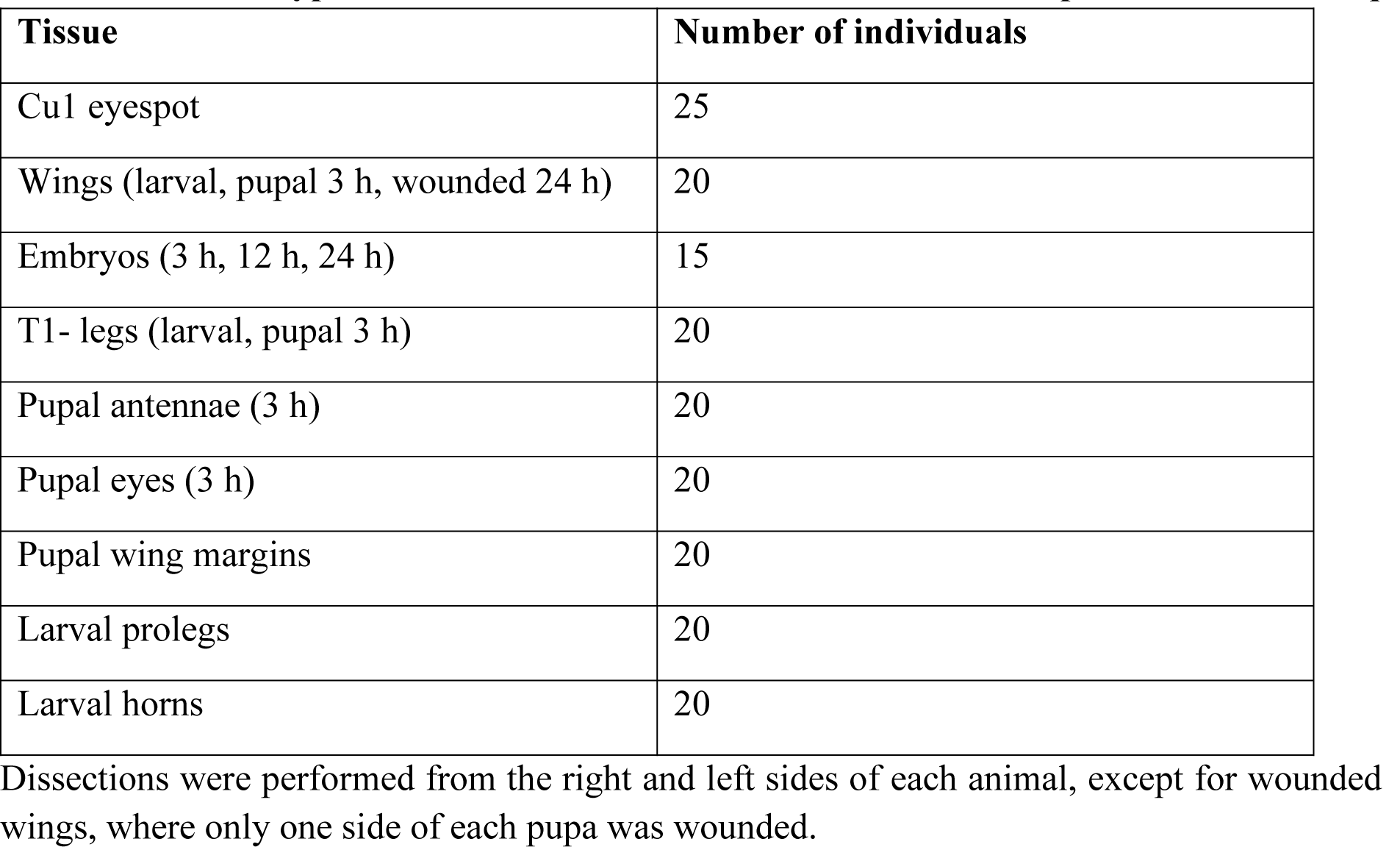
Tissues types and numbers of individuals used for each replicate in ATAC-seq

| **Tissue** | **Number of individuals** |
| --- | --- |
| Cu1 eyespot | 25 |
| Wings (larval, pupal 3 h, wounded 24 h) | 20 |
| Embryos (3 h, 12 h, 24 h) | 15 |
| T1- legs (larval, pupal 3 h) | 20 |
| Pupal antennae (3 h) | 20 |
| Pupal eyes (3 h) | 20 |
| Pupal wing margins | 20 |
| Larval prolegs | 20 |
| Larval horns | 20 |

Dissections were performed from the right and left sides of each animal, except for wounded wings, where only one side of each pupa was wounded.

**Table S8:** ATAC-Seq reads. Read depth and FRiP score

| **Group** | **Sample** | **Reads mapped to genome** | **Reads mapped to peaks** | **FRiP score** |
| --- | --- | --- | --- | --- |
| **Embryos (4 h)** | A1 | 7929162 | 7069208 | 0.892 |
|  | A2 | 8722848 | 7650722 | 0.878 |
|  | A3 | 7590222 | 6645944 | 0.876 |
|  | A4 | 8079972 | 7027172 | 0.87 |
| **Embryos (12 h)** | B1 | 28845464 | 24414192 | 0.847 |
|  | B2 | 21022574 | 17806982 | 0.848 |
|  | B3 | 26855374 | 22789314 | 0.849 |
|  | B4 | 24438812 | 20817352 | 0.852 |
| **Embryos (24 h)** | C1 | 23947046 | 20470934 | 0.855 |
|  | C2 | 19317116 | 16720430 | 0.866 |
|  | C3 | 17459032 | 15060032 | 0.863 |
|  | C4 | 23499084 | 20482478 | 0.872 |
| **Larval prolegs** | D1 | 17098004 | 14066208 | 0.823 |
|  | D2 | 16925178 | 13848684 | 0.819 |
|  | D3 | 18225832 | 14975016 | 0.822 |
| **Larval forewings** | E1 | 22238552 | 19258772 | 0.867 |
|  | E2 | 22896714 | 20094916 | 0.878 |
|  | E4 | 22130896 | 19302416 | 0.873 |
| **Pupal forewings (3 h)** | F1 | 32928940 | 28260602 | 0.859 |
|  | F2 | 25811542 | 22215800 | 0.861 |
|  | F3 | 30779202 | 26438266 | 0.859 |
|  | F4 | 30407764 | 26060672 | 0.858 |
| **Pupal antennae (3 h)** | G1 | 39990330 | 33025110 | 0.826 |
|  | G2 | 26276830 | 21934216 | 0.835 |
|  | G3 | 35808400 | 29831484 | 0.834 |
| **Pupal T1-legs (3 h)** | H1 | 35962956 | 29719496 | 0.827 |
|  | H3 | 30691458 | 25407180 | 0.828 |
|  | H4 | 32904694 | 27491442 | 0.836 |
| **Pupal eyes (3 h)** | K1 | 27369070 | 23057554 | 0.843 |
|  | K2 | 25115380 | 21124918 | 0.842 |
|  | K3 | 23566544 | 19761176 | 0.839 |
| **Larval T1-legs** | L1 | 24453864 | 20554604 | 0.841 |
|  | L2 | 28109100 | 22717706 | 0.809 |
|  | L3 | 27081264 | 22121078 | 0.817 |
|  | L4 | 22582662 | 18430048 | 0.817 |
| **Pupal eyespots (3 h)** | M1 | 23591594 | 19145932 | 0.812 |
|  | M2 | 24294062 | 20029744 | 0.825 |
| **Larval horns** | O1 | 21485294 | 17388048 | 0.81 |
|  | O2 | 20946732 | 16941090 | 0.809 |
|  | O3 | 19939442 | 16082748 | 0.807 |
|  | O4 | 27568916 | 21767434 | 0.79 |
| **Pupal forewing margins (3 h)** | Q1 | 79547324 | 65788012 | 0.828 |
|  | Q2 | 31692538 | 26044216 | 0.822 |
|  | Q3 | 24654044 | 20246276 | 0.822 |
|  | Q4 | 25965642 | 21625304 | 0.833 |
| **Wounded**<br>**pupal wings**<br>**(24 h)** | W1 | 30988206 | 26493042 | 0.855 |
|  | W2 | 36814878 | 31509656 | 0.856 |
|  | W3 | 49790726 | 43261278 | 0.869 |
|  | W4 | 27833492 | 23647830 | 0.85 |

## References

1. W. J. Glassford, W. C. Johnson, N. R. Dall, S. J. Smith, Y. Liu, W. Boll, M. Noll, M. Rebeiz, Co-option of an Ancestral Hox-Regulated Network Underlies a Recently Evolved Morphological Novelty. Dev. Cell. 34, 520–531 (2015).

2. S. B. Carroll, J. Gates, D. N. Keys, S. W. Paddock, G. E. Panganiban, J. E. Selegue, J. a Williams, Pattern formation and eyespot determination in butterfly wings. Science. 265, 109–14 (1994).

3. S. V Saenko, V. French, P. M. Brakefield, P. Beldade, Conserved developmental processes and the formation of evolutionary novelties: examples from butterfly wings. Philos. Trans. R. Soc. Lond. B. Biol. Sci. 363, 1549–55 (2008).

4. L. I. Held, Rethinking Butterfly Eyespots. Evol. Biol. 40, 158–168 (2013).

5. A. Monteiro, G. Glaser, S. Stockslager, N. Glansdorp, D. Ramos, Comparative insights into questions of lepidopteran wing pattern homology. BMC Dev. Biol. 6, 1–13 (2006).

6. H. Connahs, S. Tlili, J. van Creij, T. Y. J. Loo, T. Das Banerjee, T. E. Saunders, A. Monteiro, Activation of butterfly eyespots by Distal-less is consistent with a reaction-diffusion process. Dev. 146, 1–12 (2019).

7. J. M. Musser, G. P. Wagner, Character trees from transcriptome data: Origin and individuation of morphological characters and the so-called “species signal.” J. Exp. Zool. Part B Mol. Dev. Evol. 324, 588–604 (2015).

8. M. I. Love, W. Huber, S. Anders, Moderated estimation of fold change and dispersion for RNA-seq data with DESeq2. Genome Biol. 15, 1–21 (2014).

9. R. Suzuki, H. Shimodaira, Pvclust: An R package for assessing the uncertainty in hierarchical clustering. Bioinformatics. 22, 1540–1542 (2006).

10. N. Özsu, A. Monteiro, Wound healing, calcium signaling, and other novel pathways are associated with the formation of butterfly eyespots. BMC Genomics. 18 (2017), doi:10.1186/s12864-017-4175-7.

11. J. D. Uhl, A. Zandvakili, B. Gebelein, A Hox Transcription Factor Collective Binds a Highly Conserved Distal-less cis-Regulatory Module to Generate Robust Transcriptional Outcomes. PLoS Genet. 12, 1–26 (2016).

12. J. T. Wagner-Bernholz, O. Wilson, G. Gibson, R. Schuh, W. J. Gehring, Identification of target genes of the homeotic gene Antennapedia by enhancer detection (Genes and Development 5 (2467-2480)). Genes Dev. 6, 328 (1992).

13. P. D. Si Dong, J. Chu, G. Panganiban, Coexpression of the homeobox genes Distal-less and homothorax determines Drosophila antennal identity. Development. 127, 209– 216 (2000).

14. J. C. Oliver, X. L. Tong, L. F. Gall, W. H. Piel, A. Monteiro, A Single Origin for Nymphalid Butterfly Eyespots Followed by Widespread Loss of Associated Gene Expression. PLoS Genet. 8 (2012), doi:10.1371/journal.pgen.1002893.

15. Y. Matsuoka, A. Monteiro, Hox genes are essential for the development of novel serial homologous eyespots on the wings of Bicyclus anynana butterflies. *Gentics*, iyaa005 (2020).

16. T. Das Banerjee, A. Monteiro, Molecular mechanisms underlying simplification of venation patterns in holometabolous insects. Dev. dev.196394 (2020), doi:10.1242/dev.196394.

17. L. Zhang, R. D. Reed, Genome editing in butterflies reveals that spalt promotes and Distal-less represses eyespot colour patterns. Nat. Commun. 7 (2016), doi:10.1038/ncomms11769.

18. C. R. Brunetti, J. E. Selegue, A. Monteiro, V. French, P. M. Brakefield, S. B. Carroll, The generation and diversification of butterfly eyespot color patterns. Curr. Biol. 11, 1578–1585 (2001).

19. S. V. Saenko, M. S. P. Marialva, P. Beldade, Involvement of the conserved Hox gene Antennapedia in the development and evolution of a novel trait. Evodevo. 2, 9 (2011).

20. G. Panganiban, Distal-less Function During Drosophila Appendage and Sense Organ Development. Dev. Dyn. 562, 554–562 (2000).

21. B. S. Emerald, S. M. Cohen, Spatial and temporal regulation of the homeotic selector gene Antennapedia is required for the establishment of leg identity in Drosophila. Dev. Biol. 267, 462–472 (2004).

22. Y. T. Lai, K. D. Deem, F. Borràs-Castells, N. Sambrani, H. Rudolf, K. Suryamohan, E. El-Sherif, M. S. Halfon, D. J. McKay, Y. Tomoyasu, Enhancer identification and activity evaluation in the red flour beetle, Tribolium castaneum. Dev. 145 (2018), doi:10.1242/dev.160663.

23. G. Panganiban, J. L. R. Rubenstein, Developmental functions of the Distal-less/Dlx homeobox genes. Development. 129, 4371–86 (2002).

24. H. S. Bruce, N. H. Patel, Knockout of crustacean leg patterning genes suggests that insect wings and body walls evolved from ancient leg segments (2020), vol. 4.

25. C. M. Clark-Hachtel, Y. Tomoyasu, Two sets of candidate crustacean wing homologues and their implication for the origin of insect wings. *Nat*. Ecol. Evol. 4, 1694–1702 (2020).

26. G. Sabarís, I. Laiker, E. Preger-Ben Noon, N. Frankel, Actors with Multiple Roles: Pleiotropic Enhancers and the Paradigm of Enhancer Modularity. Trends Genet. 35, 423–433 (2019).

27. B. Prud’homme, N. Gompel, S. B. Carroll, Emerging principles of regulatory evolution. Light Evol. 1, 109–127 (2007).

28. A. Monteiro, O. Podlaha, Wings, horns, and butterfly eyespots: how do complex traits evolve? PLoS Biol. 7, e37 (2009).

29. H. Li, R. Durbin, Fast and accurate short read alignment with Burrows-Wheeler transform. Bioinformatics. 25, 1754–1760 (2009).

30. H. Li, B. Handsaker, A. Wysoker, T. Fennell, J. Ruan, N. Homer, G. Marth, G. Abecasis, R. Durbin, The Sequence Alignment/Map format and SAMtools. Bioinformatics. 25, 2078–2079 (2009).

31. A. R. Quinlan, I. M. Hall, BEDTools: A flexible suite of utilities for comparing genomic features. Bioinformatics. 26, 841–842 (2010).

32. Y. Zhang, T. Liu, C. A. Meyer, J. Eeckhoute, D. S. Johnson, B. E. Bernstein, C. Nussbaum, R. M. Myers, M. Brown, W. Li, X. S. Shirley, Model-based analysis of ChIP-Seq (MACS). Genome Biol. 9 (2008), doi:10.1186/gb-2008-9-9-r137.

33. Y. Naito, K. Hino, H. Bono, K. Ui-Tei, CRISPRdirect: Software for designing CRISPR/Cas guide RNA with reduced off-target sites. Bioinformatics. 31, 1120–1123 (2015).

34. K. N. Eckermann, H. M. M. Ahmed, M. KaramiNejadRanjbar, S. Dippel, C. E. Ogaugwu, P. Kitzmann, M. D. Isah, E. A. Wimmer, Hyperactive piggyBac transposase improves transformation efficiency in diverse insect species. Insect Biochem. Mol. Biol. 98, 16–24 (2018).

35. C. J. Evans, J. M. Olson, K. T. Ngo, E. Kim, N. E. Lee, E. Kuoy, A. N. Patananan, D. Sitz, P. T. Tran, M. T. Do, K. Yackle, A. Cespedes, V. Hartenstein, G. B. Call, U. Banerjee, G-TRACE: Rapid Gal4-based cell lineage analysis in Drosophila. Nat. Methods. 6, 603–605 (2009).

36. B. Bushnell, BBMap: A Fast, Accurate, Splice-Aware Aligner. United States N. p (2014), (available at https://www.osti.gov/biblio/1241166-bbmap-fast-accurate-splice-aware-aligner).

37. E. Kopylova, L. Noé, H. Touzet, SortMeRNA: Fast and accurate filtering of ribosomal RNAs in metatranscriptomic data. Bioinformatics. 28, 3211–3217 (2012).

38. M. Pertea, D. Kim, G. M. Pertea, J. T. Leek, S. L. Salzberg, Transcript-level expression analysis of RNA-seq experiments with HISAT, StringTie and Ballgown. Nat. Protoc. 11, 1650–1667 (2016).

39. M. S. Campbel, C. Holt, B. Moore, M. Yandell, Genome Annotation and Curation Using MAKER and MAKER-P (2008), vol. 48.

40. B. J. Haas, A. Papanicolaou, M. Yassour, M. Grabherr, D. Philip, J. Bowden, M. B. Couger, D. Eccles, B. Li, M. D. Macmanes, M. Ott, J. Orvis, N. Pochet, F. Strozzi, N. Weeks, R. Westerman, T. William, C. N. Dewey, R. Henschel, R. D. Leduc, N. Friedman, A. Regev, De novo transcript sequence recostruction from RNA-Seq: reference generation and analysis with Trinity (2013), vol. 8.

41. C. R. Fisher, J. L. Wegrzyn, E. L. Jockusch, Co-option of wing-patterning genes underlies the evolution of the treehopper helmet. *Nat*. Ecol. Evol. 4, 250–260 (2020).

42. R Core Team, R: A language and environment for statistical computing (2020).

43. M. R. Corces, A. E. Trevino, E. G. Hamilton, P. G. Greenside, N. A. Sinnott- Armstrong, S. Vesuna, A. T. Satpathy, A. J. Rubin, K. S. Montine, B. Wu, A. Kathiria, S. W. Cho, M. R. Mumbach, A. C. Carter, M. Kasowski, L. A. Orloff, V. I. Risca, A. Kundaje, P. A. Khavari, T. J. Montine, W. J. Greenleaf, H. Y. Chang, An improved ATAC-seq protocol reduces background and enables interrogation of frozen tissues. Nat. Methods. 14, 959–962 (2017).

44. A. P. Boyle, J. Guinney, G. E. Crawford, T. S. Furey, F-Seq: A feature density estimator for high-throughput sequence tags. Bioinformatics. 24, 2537–2538 (2008).

45. Y. Liao, G. K. Smyth, W. Shi, FeatureCounts: An efficient general purpose program for assigning sequence reads to genomic features. Bioinformatics. 30, 923–930 (2014).

46. F. Ramírez, F. Dündar, S. Diehl, B. A. Grüning, T. Manke, DeepTools: A flexible platform for exploring deep-sequencing data. Nucleic Acids Res. 42, 187–191 (2014).

47. N. C. Durand, M. S. Shamim, I. Machol, S. S. P. Rao, M. H. Huntley, E. S. Lander, E. L. Aiden, A. Mathematics, Juicer provides a one-click system for analyzing loop-resolution Hi-C experiments. Cell Syst. 3, 95–98 (2018).

48. J. Ray, P. R. Munn, A. Vihervaara, J. J. Lewis, A. Ozer, C. G. Danko, J. T. Lis, Chromatin conformation remains stable upon extensive transcriptional changes driven by heat shock. Proc. Natl. Acad. Sci. U. S. A. 116, 19431–19439 (2019).

49. R. W. Nowell, B. Elsworth, V. Oostra, B. J. Zwaan, C. W. Wheat, M. Saastamoinen, I. J. Saccheri, A. E. van’t Hof, B. R. Wasik, H. Connahs, M. L. Aslam, S. Kumar, R. J. Challis, A. Monteiro, P. M. Brakefield, M. Blaxter, A high-coverage draft genome of the mycalesine butterfly Bicyclus anynana. Gigascience. 6, 1–7 (2017).

50. P. Beldade, S. V. Saenko, N. Pul, A. D. Long, A gene-based linkage map for Bicyclus anynana butterflies allows for a comprehensive analysis of synteny with the lepidopteran reference genome. PLoS Genet. 5 (2009), doi:10.1371/journal.pgen.1000366.

51. J. Catchen, A. Amores, S. Bassham, G3&58; Genes|Genomes|Genetics, in press, doi:10.1534/g3.120.401485.

52. F. A. Simão, R. M. Waterhouse, P. Ioannidis, E. V. Kriventseva, E. M. Zdobnov, BUSCO: Assessing genome assembly and annotation completeness with single-copy orthologs. Bioinformatics. 31, 3210–3212 (2015).

53. P. Jones, D. Binns, H. Y. Chang, M. Fraser, W. Li, C. McAnulla, H. McWilliam, J. Maslen, A. Mitchell, G. Nuka, S. Pesseat, A. F. Quinn, A. Sangrador-Vegas, M. Scheremetjew, S. Y. Yong, R. Lopez, S. Hunter, InterProScan 5: Genome-scale protein function classification. Bioinformatics. 30, 1236–1240 (2014).

54. J. Dainat, AGAT: Another Gff Analysis Toolkit to handle annotations in any GTF/GFF format.(Version v0.4.0) (2020), doi:10.5281/ZENODO.4205393.

55. B. Buchfink, C. Xie, D. H. Huson, Fast and sensitive protein alignment using DIAMOND. Nat. Methods. 12, 59–60 (2014).

56. S. Götz, J. M. García-Gómez, J. Terol, T. D. Williams, S. H. Nagaraj, M. J. Nueda, M. Robles, M. Talón, J. Dopazo, A. Conesa, High-throughput functional annotation and data mining with the Blast2GO suite. Nucleic Acids Res. 36, 3420–3435 (2008).

